# High stakes slow responding, but do not help overcome Pavlovian biases in humans

**DOI:** 10.1101/2023.12.26.573351

**Authors:** Johannes Algermissen, Hanneke E. M. den Ouden

## Abstract

“Pavlovian” or “motivational” biases are the phenomenon that the valence of prospective outcomes modulates action invigoration: the prospect of reward invigorates actions, while the prospect of punishment suppresses actions. Effects of the valence of prospective outcomes are well established, but it remains unclear how the magnitude of outcomes (“stake magnitude”) modulates these biases. In this pre-registered study (*N* = 55), we manipulated stake magnitude (high vs. low) in an orthogonalized Motivational Go/NoGo Task. We tested whether higher stakes (a) strengthen biases or (b) elicit cognitive control recruitment, enhancing the suppression of biases in motivationally incongruent conditions. Confirmatory tests showed that high stakes slowed down responding, especially in motivationally incongruent conditions. However, high stakes did not affect whether a response was made or not, and did not change the magnitude of Pavlovian biases. Reinforcement-learning drift-diffusion models (RL-DDMs) fit to the data suggested that response slowing was best captured by stakes prolonging the non-decision time. There was no effect of the stakes on the response threshold (as in typical speed-accuracy tradeoffs). In sum, these results suggest that high stakes slow down responses without affecting the expression of Pavlovian biases in behavior. We speculate that this slowing under high stakes might reflect heightened cognitive control, which is however ineffectively used, or reflect positive conditioned suppression, i.e., the interference between goal-directed and consummatory behaviors, a phenomenon previously observed in rodents that might also exist in humans. Pavlovian biases and slowing under high stakes may arise in parallel to each other.

## High stakes slow responding, but do not help overcome Pavlovian biases in humans

The behavior of humans and other animals reflects the interplay of multiple, partly independent decision-making systems (Collins and Cockburn 2020; Daw et al. 2005; Dickinson and Balleine 1994; Metcalfe and Mischel 1999; Shiffrin and Schneider 1977; Strack and Deutsch 2004). One such system is the Pavlovian system which rigidly triggers response invigoration to the prospect of reward and response inhibition to the threat of punishment (Boureau and Dayan 2011; O’Doherty et al. 2017; Guitart-Masip et al. 2014). Its actions are visible in the form or “Pavlovian” or “motivational” biases, which have been proposed to underlie many seemingly maladaptive behaviors in humans and other animals (Dayan et al. 2006).

Learning about the positive/negative consequences of our actions has been proposed to happen via two distinct learning systems (Dayan and Huys 2009; Boureau and Dayan 2011): the instrumental system learns the relationship between actions and ensuing outcomes, while the Pavlovian system learns which outcomes can be expected in a particular situation irrespective of the actions taken. In other words, the Pavlovian system tracks outcomes that an agent does not have to actively contribute to. Still, animals and humans show so-called unconditioned responses to the outcomes themselves, such as approach to appetitive objects and avoidance/withdrawal from aversive outcomes. With learning, the Pavlovian system transfers these responses to neutral cues that predict these outcomes, becoming conditioned responses. These conditioned responses may interfere with instrumental behavior when an agent has to select the right action in order to obtain a desired outcome (Boakes et al. 1978). In such cases, the instrumentally learned action that should be selected for a certain outcome and the Pavlovian response involuntarily triggered in anticipation of that outcome can work in concert, e.g. when obtaining a reward involves approaching it, but can also clash, e.g. when obtaining the reward requires passive waiting. When instrumental and Pavlovian responses conflict, they become visible in the form of Pavlovian biases, with the Pavlovian system involuntarily triggering responses that prevent the instrumental system from reaching its goals (Breland and Breland 1961; Hershberger 1986).

Pavlovian mechanisms might explain seemingly “irrational” behaviors in animals. Sometimes, irrelevant, but reward-predictive cues can facilitate instrumental approach behavior on a focal task (Estes 1943, 1948; Rescorla and Solomon 1967; LoLordo et al. 1974; Schwartz 1976; Lovibond 1983). In other situations, reward-predictive cues can distract an animal from a focal task, called “sign-tracking” behavior (Jenkins and Moore 1973; Hearst and Jenkins 1974). Sign-tracking is interpreted as rigid, inflexible behavior that is overly controlled by stimuli signaling the chance for rewards, but less influenced by whether a reward is eventually obtained or not. It likely reflects conditioned approach responses and thus an instance of Pavlovian biases preventing instrumental behavior towards the goal. Sign-tracking animals have been proposed as a model to study human drug addiction, with the neural mechanisms underlying sign-tracking potentially explaining how drug-related stimuli can induce strong drug cravings (Flagel et al. 2009; Saunders and Robinson 2010). Hence, sign-tracking might be a phenomenon shared across species, including humans (Colaizzi et al. 2020; Garofalo and di Pellegrino 2015) and might contribute to the etiology and maintenance of drug abuse (Flagel and Robinson 2017). A better understanding of when Pavlovian biases occur and how they interact with other systems regulating behavior promises insights into the development and maintenance of psychiatry conditions such as alcohol or drug abuse (Chen et al. 2022b; Schad et al. 2020).

While the occurrence of Pavlovian biases seems well established, it is unclear whether and how the magnitude of anticipated outcomes (“stakes”) influence the size of these biases. Conditioned responses are typically sensitive to the amount of reward (or shock) predicted: if multiple stimuli predict one unit of food each, an animal will expect their sum, and will show stronger anticipatory responding (summation effect) (Khallad and Moore 1996; Rescorla 1999; Kehoe and White 2004). However, if only one unit of food is provided, the response to each individual stimulus decreases over time (over- expectation effect) (Rescorla 1970, 1999). Similarly, humans show faster and more vigorous responses in anticipation of larger rewards, even if speed or vigor are irrelevant for obtaining the reward (Knutson et al. 2005; Pessiglione et al. 2007; Bijleveld et al. 2010). Neural network models of the basal ganglia also predict that larger stakes lead to larger Pavlovian biases: reward-predictive stimuli elicit positive prediction errors in the form of dopamine bursts, which sensitize the direct pathway of the basal ganglia and thus facilitate the invigoration of behavior (Frank 2005; Collins and Frank 2014). Stimuli predicting larger rewards should elicit larger prediction errors, larger dopamine bursts, and thus even higher invigoration. Vice versa, punishment-predictive stimuli elicit negative prediction errors and dopamine dips, which sensitize the indirect pathway and thus facilitate the suppression of behavior, an effect expected to be stronger for stimuli predicting larger punishments. In sum, both previous behavioral evidence on conditioned responses as well as computational network models of the basal ganglia predict that stimuli signaling larger rewards/punishments should elicit stronger Pavlovian biases.

Evidence that the strength of Pavlovian biases varies with stake magnitude has been mixed so far. A few studies using Pavlovian-to-Instrumental Transfer (PIT) tasks, in which task-irrelevant cues associated with rewards/punishments are presented in the background, have observed slight increases in response rates and somewhat faster reaction times for higher rewards (Algermissen and den Ouden 2023; Schad et al. 2020) as well as decreased response rates and slower reaction times for larger punishments (Geurts et al. 2013a, 2013b). However, many other studies have not observed such modulations (Chen et al. 2022a, 2023; Garbusow et al. 2019, 2016; Sommer et al. 2017, 2020). Similarly, some studies have found PIT effects to become weaker when rewards are devalued through satiation or pairing with aversive cues (Colwill and Rescorla 1990; Corbit et al. 2007; Allman et al. 2010), but other studies have failed to observe such devaluation effects (Rescorla 1994; Holland 2004; Watson et al. 2014; Pool et al. 2019). Beyond the PIT task, other paradigms that varied the reward on offer, specifically versions of the monetary incentive delay task, have observed faster reaction times to larger rewards (Knutson et al. 2005; Luo et al. 2009). A study using a virtual predation game found slower reaction times under larger threats (Bach 2015). However, in these studies, it remained unclear whether reward-induced invigoration/punishment-induced slowing followed from automatic, Pavlovian effects or rather participants’ deliberate strategies, reflecting their beliefs about which behavior was conducive to reward attainment/punishment avoidance (Mahlberg et al. 2021; Westbrook et al. 2021). To disentangle automatic from strategic effects, there must be task conditions that incentivize the suppression of Pavlovian biases—a unique feature of the Motivational Go/NoGo Task. In this task, individuals learn through trial-and-error to perform or withhold responses to cues in order to gain points (Win cues) or avoid loss of points (Avoid cues) (Fig. 1A-C). Humans show more (and faster) Go responses to Win than Avoid cues, reflecting the influence of Pavlovian biases on instrumental responding (Guitart-Masip et al. 2012, 2014; Swart et al. 2017). These biases are typically much stronger than in PIT tasks. Importantly, in this task, the correct action required for positive outcomes is orthogonal to the bias-triggered action, such that participants can follow their bias on half of the trials, but have to inhibit it on the other half of the trials in order to perform the correct response.

**Figure 1.**
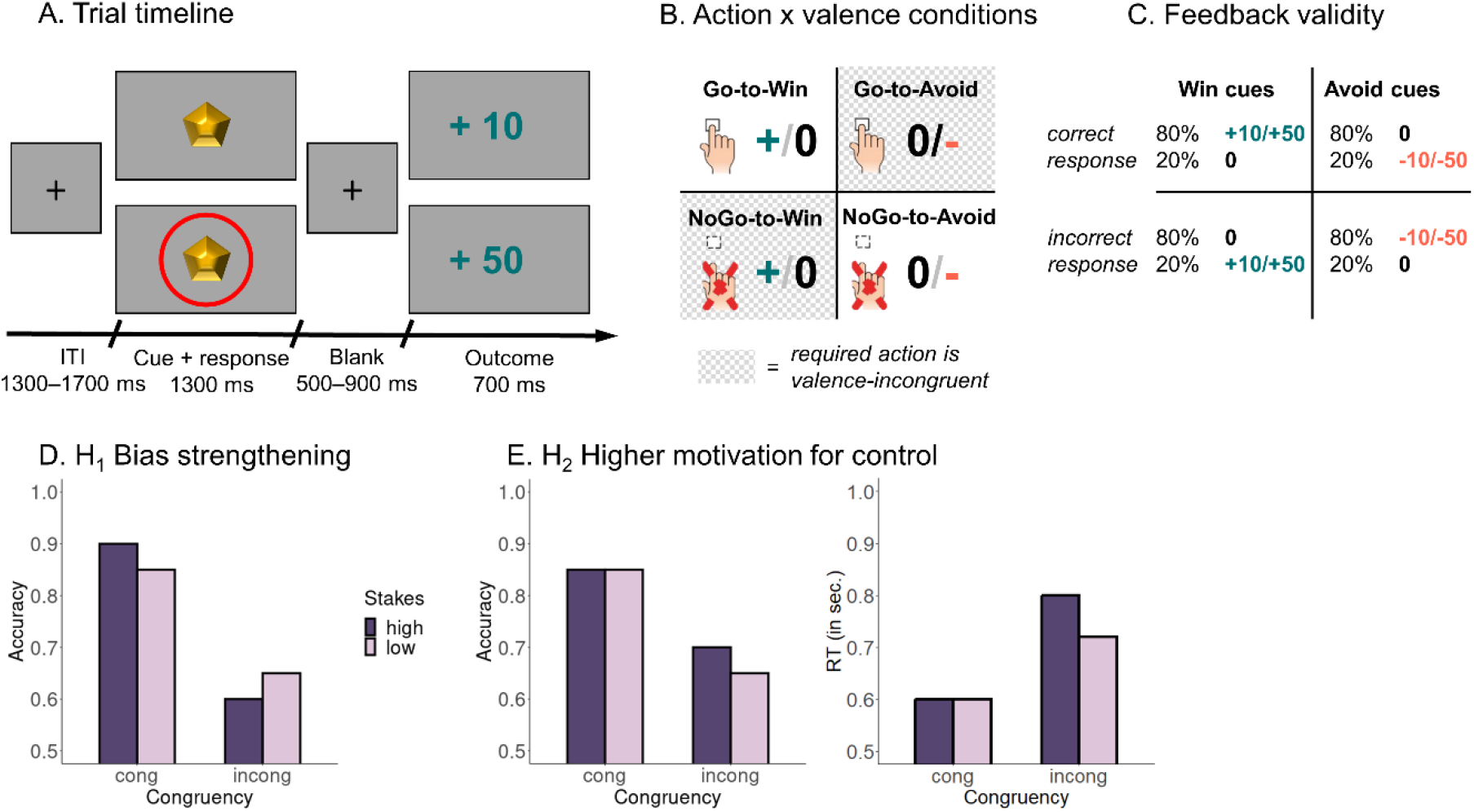
Task and behavioral predictions. **A.** Time course of each trial. Participant see one of four cues (“gems”) and have to decide whether to respond to it with a button press (“Go”) or not (“NoGo”). On half of the trials, the cue is surrounded by a red circle, indicating that stakes are five times as high and points gained/lost in this trial will be multiplied with 5. After a variable interval, participants receive an outcome (increase in points, no change, or decrease in points). **B.** Task conditions. Half of the cues are “Win” cues for which points can be gained (or no change in the point score occurs), while the other half are “Avoid” cues for which points can be lost (or no change in the point score occurs). Orthogonal to cue valence is the correct action required for each cue, which is either Go or NoGo. **C.** Feedback given cue valence and response accuracy. For Win cues, correct responses mostly lead to an increase in points (+10 or +50, depending on whether the trial was high or low stakes), but occasionally lead to no change in score (0). For Avoid cues, correct responses mostly lead to no change in score (0), while occasionally lead to a loss of points (-10 or -50, depending on whether the trial was high or low stakes). For incorrect responses, probabilities are reversed. **D.** Prediction from a “bias strengthening” hypothesis. High stakes strengthen biases, leading to higher accuracy for bias-congruent cues (for which required action and valence match), but lower accuracy for bias-incongruent cues. **E.** Prediction from the “expected value of control” theory. High stakes motivate cognitive control, which inhibits biases when they are incongruent with the required action, leading to higher accuracy selectively for bias-incongruent cues (for which the bias-triggered response has to be inhibited). At the same time, cognitive control recruitment takes time, leading to slower reaction times on incongruent trials, and this slowing is stronger under high stakes when more control is recruited.

While Pavlovian biases might lead to adaptive behavior in a number of situations, they become most apparent in situations in which they conflict with optimal behavior: sometimes, agents have to wait to secure a reward, e.g., in situations akin to the Marshmallow Test (Mischel and Ebbesen 1970), or they have to take active steps to prevent or fight a threat, e.g., in exposure therapy to treat arachnophobia. In such circumstances, agents have to suppress Pavlovian biases, a requirement animals usually struggle with (Breland and Breland 1961; Hershberger 1986) and even humans only imperfectly master (Cavanagh et al. 2013; Swart et al. 2018). The ability to suppress automatic, unwanted action tendencies is regarded to require cognitive control (Cohen 2017). For several decades, cognitive control has been seen as a limited resource or ability that can fail, leading to action slips and undesired behavior (Hofmann et al. 2009). In contrast, more recent perspectives, most notably the *expected value of control theory* (EVC) (Shenhav et al. 2013; Lieder et al. 2018) have suggested that cognitive control is not inherently limited, but follows from a cost-benefit trade-off that weighs the benefits of exerting additional control against the costs of doing so. In line with this idea, a number of studies using conflict tasks, such as the Stroop, Simon, or Flanker task, have shown that congruency effects—taken to reflect cognitive control limitations—become smaller when participants are offered financial incentives for recruiting control (Boehler et al. 2012; Chiew and Braver 2014; Dixon and Christoff 2012; Fröber and Dreisbach 2016; Krebs et al. 2010). From this perspective, higher stakes should motivate an agent to exert additional cognitive control in order to suppress biases in situations in which those are maladaptive. The EVC theory thus makes predictions directly opposite to the above-described case of high stakes strengthening biases: EVC predicts that higher stakes should lead to more cognitive control recruitment and thus weaker biases. Preliminary evidence for this claim comes from work showing that Pavlovian biases are stronger in uncontrollable situations (i.e., when correct and incorrect responses are rewarded at the same rate) in which there is no benefit to instrumentally learning action-outcome-contingencies (Dorfman and Gershman 2019). However, this work only considered the relative difference in outcomes for correct and incorrect responses and distinguished controllable from uncontrollable environments. In contrast, EVC predicts that, across a range of controllable environments (i.e., when learning action-outcome-contingencies is beneficial), the absolute magnitude of outcomes should modulate the amount of control exerted and thus the strength of Pavlovian biases.

In this study, we directly tested these two opposing predictions against each other. We collected data from 55 participants performing the motivational Go/NoGo Task in which the magnitude of stakes (high or low) was manipulated on a trial-by-trial basis. Irrespective of the stakes, we expected a congruency effect, i.e., higher accuracy in conditions in which the required response matches the bias- triggered response (Go-to-Win, NoGo-to-Avoid) compared to conditions in which the required response and bias mismatch (Go-Avoid, NoGo-to-Win). Following the first hypothesis that higher stakes drive stronger Pavlovian biases, we predicted an interaction between congruency and the stakes magnitude, with a stronger congruency effect and higher performance on congruent, but lower performance on incongruent trials under high compared to low stakes (Fig. 1D). In contrast, following the EVC theory, we predicted an interaction effect in the opposite direction, with a weaker congruency effect (reflecting cognitive control recruitment) and selectively higher performance on incongruent trials under high compared to low stakes (Fig. 1E). In addition, cognitive control recruitment is assumed to take time; hence, reaction times should be slower on incongruent trials, but particularly slow under high stakes when more control is recruited. Thus, we also predicted an interaction effect between congruency and stakes on reaction times, with a stronger congruency effect under high stakes (Fig. 1E).

## Results

### Approach

Fifty-five participants (54 included in analyses) played a adapted version of the Motivational Go/NoGo Learning Task (Swart et al. 2017). This task required them to learn from trial-and-error whether to perform a Go response (button press) or NoGo response (no response) to various cues (Fig. 1A). Half of the cues required a Go response (Go cues), the other half a NoGo response (NoGo cues; Fig. 1B). Orthogonal to the required action, half of cues offered the chance to win points for correct responses (Win cues; no change in points for incorrect responses), while the other half bore the chance to lose points for incorrect responses (Avoid cues; no change in points for correct responses). Participants typically show a Pavlovian bias in this task, with more Go responses and faster RTs for Win than Avoid cues. Feedback was probabilistic, with correct responses leading to desired outcomes on 80% of trials (win for Win cues, no change for Avoid cues), but undesired outcomes on the remaining 20% of trials (no change for Win cues, loss for Avoid cues; probabilities were reversed for incorrect responses; Fig. 1C). Orthogonal to both the required action and the valence (Win/ Avoid) of cues, we varied the stake magnitude: On half of the trials, the cue was surrounded by a red circle, signaling the chance to win/lose 50 points (instead of 10 points) for correct/incorrect responses.

### Sanity checks: Learning and Pavlovian biases

As a sanity check and in order to compare the results from this study to previous studies (Algermissen et al. 2022; Swart et al. 2018, 2017), we fitted a mixed-effects logistic regression with responses (Go/NoGo) as dependent variable as well as required action (Go/NoGo) and valence (Win/Avoid) and independent variables (see Supplementary Material S01 for an overview of all regression results; see Supplementary Material S04 for means and standard deviations per condition). In line with previous findings, participants made significantly more Go responses to Go cues than NoGo cues (required action), *b* = 1.441, 95%-CI [1.252, 1.630], χ^2^(1) = 87.873, *p* < .001, indicating that they learned the task. They also showed significantly more Go responses to Win than Avoid cues (cue valence), *b* = 0.750, 95%-CI [0.609, 0.889], χ^2^(1) = 59.587, *p* < .001, reflecting a Pavlovian bias (Fig. 2A–C). There was no evidence for the Pavlovian bias being stronger for either Go or NoGo cues (required action x valence), *b* = 0.019, 95%-CI [-0.100, 0.137], χ^2^(1) = 0.093, *p* = .760.

**Figure 2.**
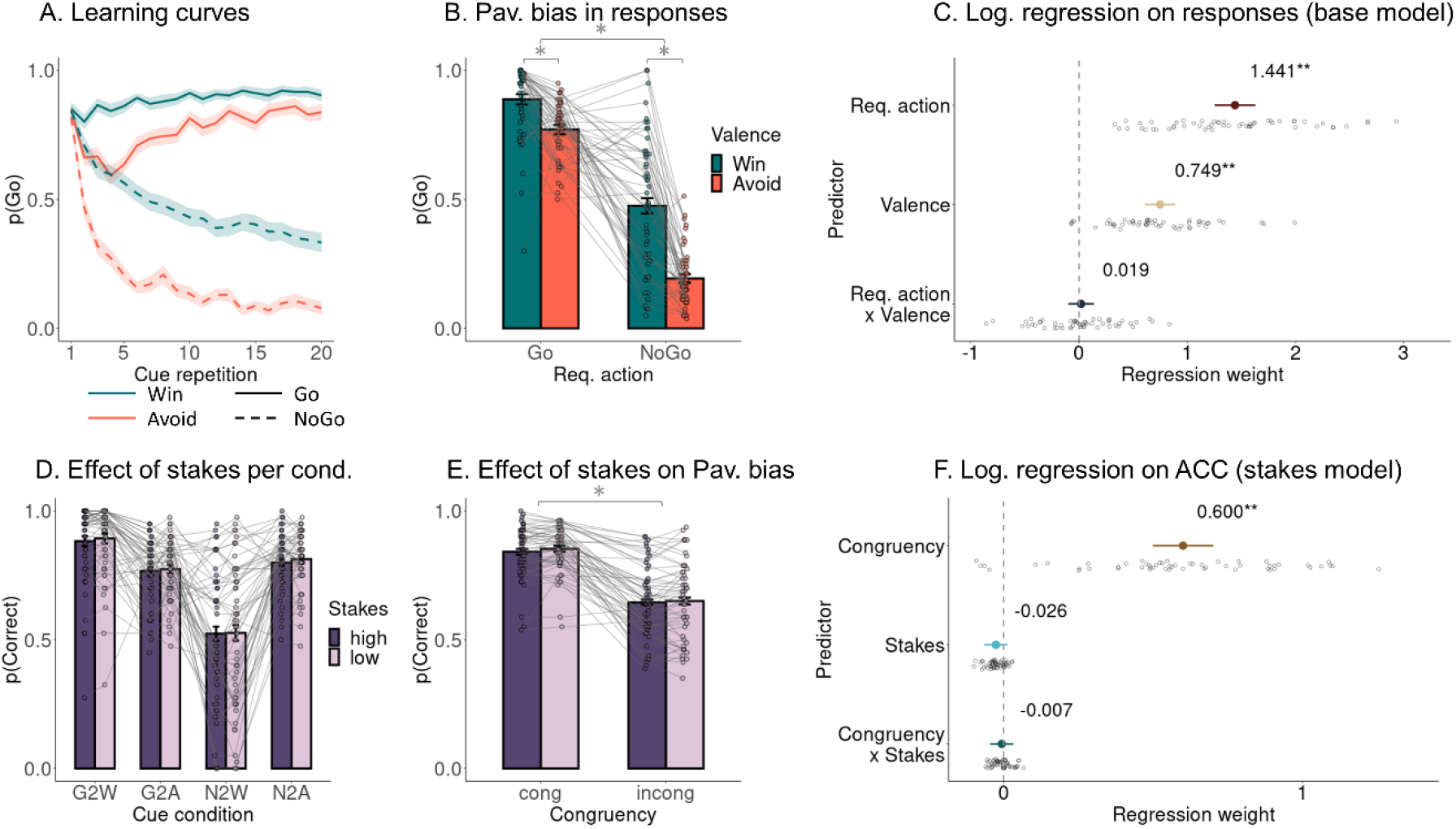
Effect on propensity of Go responses. **A.** Learning curves per cue condition. **B.** Proportion of Go responses per cue condition (individual dots are individual participant means). Participants show more Go responses to Go than NoGo cues (indicative of learning the task) and more Go responses to Win cues than Avoid cues (indicative of Pavlovian biases). **C.** Group-level (colored dot, 95%-CI) and individual-participant (grey dots) regression coefficients from a mixed-effects logistic regression of responses on required action, cue valence, and their interaction. **D.** Accuracy per cue condition and stakes condition. Accuracy is higher for congruent than incongruent cues, but there is no effect of stakes on responses for any cue condition. **E.** Accuracy per valence-action congruency and stakes condition. Accuracy is higher for congruent than incongruent conditions, but this congruency effect is not modulated by stakes. **F.** Group-level and individual-participant regression coefficients from a mixed-effects logistic regression of responses on congruency, stakes, and their interaction.

Next, we performed a similar mixed-effects linear regression with reaction times (RTs) as dependent variable. Note that RTs were naturally only available for (correct and incorrect) Go responses. Participants showed significantly faster (correct) responses to Go cues than (incorrect) responses to NoGo cues (required action), *b* = -0.109, 95%-CI [-0.145, -0.073], χ^2^(1) = 27.494, *p* < .001, and significantly faster responses to Win than Avoid cues (cue valence), *b* = -0.191, 95%-CI [-0.227, - 0.155], χ^2^(1) = 59.204, *p* < .001, again reflecting the Pavlovian bias (Fig. 3A–C). The cue valence effect (Pavlovian bias) on RTs was slightly stronger for (correct) response to Go cues than (incorrect) responses to NoGo cues (required action x cue valence), *b* = -0.032, 95%-CI [-0.061, -0.003], χ^2^(1) = 4.384, *p* = .036. The strength of the Pavlovian bias (both in responses and RTs) was neither correlated with working memory span, nor impulsivity, nor neuroticism (Supplementary Material S05). In sum, participants learned the task and exhibited a Pavlovian bias in both responses and RTs.

**Figure 3.**
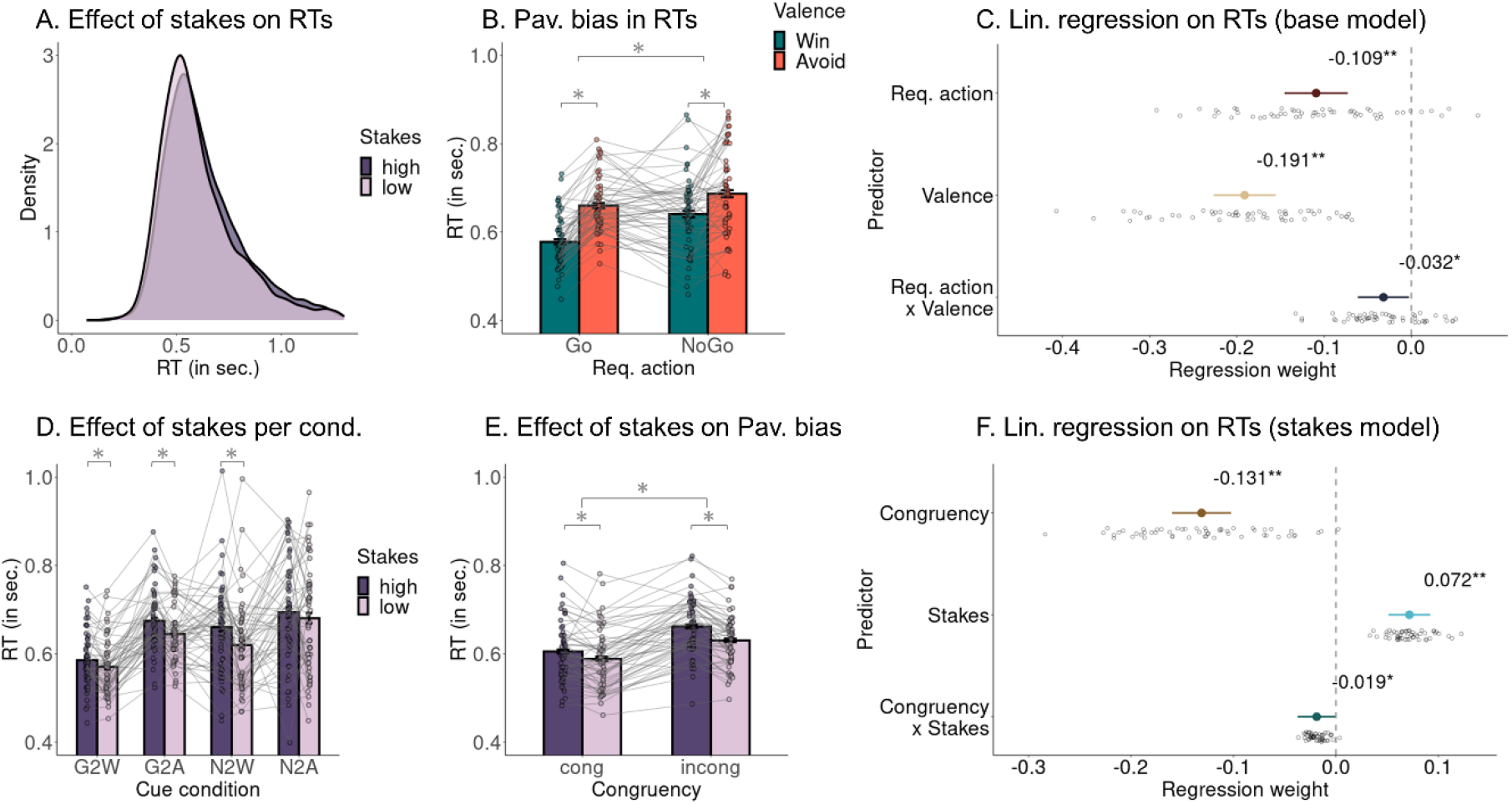
Effect on propensity of reaction times (RTs). **A.** Distribution of RTs for high and low stakes. RTs are slower under high stakes. **B.** RTs per cue condition. Participants show faster RTs for (correct) Go responses to Go cues than (incorrect) Go responses to NoGo cues. Also, they show faster RTs to Win cues than Avoid cues, indicative of Pavlovian biases in the form of invigoration under reward prospects and inhibition under threat of punishment. **C.** Group-level (colored dot, 95%-CI) and individual-participant (grey dots) regression coefficients from a mixed-effects linear regression of RTs on required action, cue valence, and their interaction. **D.** RTs per cue condition and stakes condition. RTs are significantly slower under high stakes in the Go-to-Win (G2W), Go-to-Avoid (G2A), and NoGo-to-Win (NG2W) conditions. **E.** RTs per valence-action congruency and stakes condition. RTs after significantly slower under high compared to low stakes. This effect is significantly stronger for incongruent than congruent cue conditions. **F.** Group-level and individual-participant regression coefficients from a mixed-effects linear regression of RTs on congruency, stakes, and their interaction.

### Confirmatory analyses: Modulation by stakes

As the first set of confirmatory, pre-registered analyses, we fitted a mixed-effects logistic regression with accuracy (correct/incorrect) as dependent variable and congruency (congruent/incongruent) and stakes (high/low) as independent variables. There was a significant main effect of congruency, *b* = 0.600, 95%-CI [0.499, 0.702], χ^2^(1) = 67.867, *p* < .001, with higher accuracy to congruent than incongruent cues, again reflecting the Pavlovian bias. However, neither the main effect of stakes, *b* = -0.026, 95%-CI [-0.065, 0.013], χ^2^(1) = 1.430, *p* = .232, nor the interaction between congruency and stakes, *b* = -0.007, 95%-CI [0.046, 0.032], χ^2^(1) = 0.094, *p* = .759, was significant (Fig. 2E, F).

Exploratory post-hoc tests for each cue condition separately did not show any effect of stakes on responses for any cue condition (Go-to-Win: *z* = -0.590, *p* = .555; Go-to-Avoid: *z* = -0.184, *p* = .854; NoGo-to-Win: *z* = -0.145, *p* = .885; NoGo-to-Avoid: *z* = -0.963, *p* = .336; Fig. 2D). In further exploratory analyses, we tested whether an effect of stakes on responses emerged (or disappeared) over time, either within the learning trajectory of a cue (cue repetition; 1–20) or across the entire task (trial number: 1–320). Neither the interaction between cue repetition and stakes, *b* = -0.002, 95%-CI [-0.039, 0.035], χ^2^(1) = 0.020, *p* = .898, nor the interaction between trial number and stakes, *b* = -0.012, 95%- CI [-0.048, 0.023], χ^2^(1) = 0.401, *p* = .527, was significant, providing no evidence for stakes influencing responses selectively at certain time points during learning or during the task. In sum, there was no evidence for stakes modulating the Pavlovian bias in participants’ responses.

As the second set of confirmatory, pre-registered analyses, we fitted a mixed-effects linear regression with RTs as dependent variable and congruency (congruent/incongruent) and stakes (high/low) as independent variables. Participants responded significantly more slowly to incongruent than congruent cues (congruency), *b* = -0.131, 95%-CI [-0.160, -0.102], χ^2^(1) = 49.546, *p* < .001. Furthermore, they responded significantly more slowly under high compared to low stakes (stakes), *b* = 0.072, 95%-CI [0.051, 0.092], χ^2^(1) = 33.702, *p* < .001 (Fig 3E, F). Finally, the interaction between congruency and stakes was significant, *b* = -0.019, 95%-CI [-0.037, -0.001], χ^2^(1) = 3.856, *p* = .049, with a stronger congruency effect under high compared to low stakes. This effect was also significant (*p* = .046) when including RTs < 300 ms (see Supplementary Material S02), but only marginally significant (*p* = .060) when adding the data of a remaining participant with not-above-chance performance (see Supplementary Material S03). The effect of stakes on RTs was correlated neither with working memory span, impulsivity, or neuroticism (Supplementary Material S05).

Exploratory (not pre-registered) post-hoc tests for each cue condition separately yielded a significant effect of stakes on RTs for three out of four cue conditions, including in particular the two incongruent conditions Go-to-Avoid and NoGo-to-Win (Go-to-Win: *z* = 2.973, *p* = .003; Go-to-Avoid: *z* = 4.528, *p* < .001; NoGo-to-Win: *z* = 4.975, *p* < .001; NoGo-to-Avoid: *z* = 1.414, *p* = .158; Fig. 3D).

In further exploratory analyses, we tested whether the effect of stakes on responses got stronger or weaker with time, either within the learning trajectory of a cue (cue repetition) or across the entire task (trial number). Neither the interaction between stakes and cue repetition, *b* = -0.012, 95%-CI [-0.030, 0.006], χ^2^(1) = 1.599, *p* = .206, nor the interaction between stakes and trial number, *b* = 0.025, 95%-CI [-0.021, 0.018], χ^2^(1) = 0.480, *p* = .489, was significant, providing no evidence for a change in the effect of stakes on RTs over time. See Supplementary Material S06 for tests for non-linear changes with time, again finding no evidence for changes in the effect of stakes over time. In sum, these results suggest that high stakes affected participants’ responses in that they overall slowed down responses. This slowing was slightly stronger for incongruent than congruent cues and appeared to be constant over time. However, stakes did not affect response accuracy nor the degree of Pavlovian bias as indexed by the decisions to make a Go or NoGo response.

### Computational Modeling of Responses and RTs (RL-DDMs)

To better understand the mechanisms by which cue valence and stakes influenced responses and RTs, we fit a series of increasingly complex reinforcement-learning drift-diffusion models (RL-DDMs) (Pedersen et al. 2017; Fontanesi et al. 2019). Drift-diffusion models are a subtype of evidence accumulation models. They take a stimulus input and transform it into a response with a certain RT.

Specifically, DDMs simulate an evolving signal that noisily accumulates (positive or negative) evidence over time with a certain speed, determined by the drift rate parameter. Once the signal hits an upper threshold, a response (e.g. “Go”) is emitted; if it instead hits a lower threshold, an alternative response (e.g. “NoGo”) is “emitted” (i.e., the decision not to respond is taken). The distance from the starting point to the response threshold is a free parameter that determines the speed-accuracy tradeoff: low thresholds lead to fast but frequently incorrect responses (due to initial random fluctuations sometimes leading to the crossing of the “incorrect” threshold), while high thresholds lead to slow but mostly correct responses (because random fluctuations become negligible compared to the evolving signal). The accumulation process can start in the middle between both thresholds or be biased towards one of them, determined by the starting point bias parameter. Lastly, sensory processing and motor execution might contribute to the RT independently of the decision process itself, and the duration of all these non-decision processes is summarized as the non-decision-time parameter.

The DDM uses the Wiener first passage time distribution to assign probability densities to each possible RT for a given response (Ratcliff 1978; Navarro and Fuss 2009), and the parameters of the model (drift rate, threshold, starting point bias, non-decision time) can be fit to observed responses and RTs. In the case of NoGo responses, no RT is observed; however, it is possible to integrate over all possible RTs of the NoGo boundary to compute the overall probability of making a NoGo response (i.e., to decide not to respond) (Gomez et al. 2007; Blurton et al. 2012). In the Motivational Go/NoGo learning task, the cues do not directly signal whether a Go or NoGo response should be performed. Instead, participants need to learn the values of Go/NoGo responses for a given cue via trial-and-error learning. These action values are updated based on received outcomes using a delta learning rule (Pedersen et al. 2017; Fontanesi et al. 2019). In RL-DDMs, the difference between the value of a Go response and the value of a NoGo response for a given cue serves as the input signal for making a decision.

Previous studies have considered different implementations of Pavlovian biases in DDMs. One possibility is to assume that cue valence modulates the starting point bias: for Win cues, the signal starts closer to the “Go” response boundary, and for Avoid cues closer to the “NoGo” response boundary.

Another possibility is that cue valence modulates the drift rate: for Win cues, evidence for “Go” is accumulated more quickly, and for Avoid cues, evidence for “NoGo” is accumulated more quickly. A past study using a similar paradigm found evidence for cue valence modulating the starting point bias rather than the drift rate (Millner et al. 2017), although evidence in that study remained mixed.

Our regression analyses described above yielded that high stakes slowed down responses. Response-slowing might reflect a speed-accuracy trade-off: under high stakes, decision thresholds might become higher, leading to higher accuracy at the cost of slower responses (Bogacz et al. 2006; Rae et al. 2014; Shevlin et al. 2022). We implemented different mechanisms of how cue valence and stakes might influence the various parameters (decision threshold, non-decision time, starting point bias, drift rate intercept) in an evidence accumulation framework and compared the fit of different, increasingly complex models.

First, we tested whether behavior was better described by an RL-DDM (M2) in which participants learned cue-specific action values over time rather than an standard DDM (M1) with a constant propensity to emit Go/NoGo responses (Fig. 4A). M2 yielded superior fit, which reflects that participants learned the task and that their learned action values influenced responses and RTs. Next, we tested whether adding a Pavlovian bias, either as a modulation of the starting point bias (M3) or as an additive bonus on the drift rate (M4) further improved model fit. Indeed, fit was best for M4 in which evidence for Go was accumulated faster for Win cues and evidence for NoGo accumulated faster for Avoid cues (Fig. 4B). Next, we assessed different mechanisms by which stake magnitude might affect responding by letting stake magnitude modulate each of the four DDM parameters (threshold, non-decision time, starting point bias, drift rate; M5–M8). Here, the best model was one in which stakes modulate the non-decision time (M6). Note that, although M6 showed clearly a superior fit to M4, group-level non-decision times for high and low stakes were not significantly different from each other (*Mdiff* = 0.012, 95%-CI [-0.017, 0.041]), suggestive of the presence of individual differences with an overall mean close to zero (Swart et al. 2017). Next, we tested whether model fit was further improved when stake magnitude could modulate two DDM parameters together (M9–M11). Specifically, a modulation of both the threshold and the drift rate is typically observed in speed-accuracy tradeoffs (M10) (Rae et al. 2014). Allowing stakes to modulate two instead of one parameter did not yield any substantial improvement in fit (M9–M11). Specifically, the model implementing a speed-accuracy tradeoff by allowing stakes to influence both the threshold and the drift rate (M10) performed worse than a model allowing stakes to influence the non-decision time (M6). Lastly, we tested whether model fit was further improved by splitting the effect of stakes into separate parameters for congruent and incongruent cues (M12). M12 was overall the best fitting model in the model comparison. Note that M12 has the same number of parameters as models M9-M11, suggesting that the superior fit of M12 is not due to a mere increase in the number of parameters, but due to the specific mechanism implemented. Also note that, although M12 with separate non-decision times under high stakes for congruent and incongruent cues outperformed M6 with a single non-decision time under high stakes, there was no group-level difference between the parameters for congruent vs. incongruent cues (*Mdiff* = -0.003, 95%- CI [-0.033, 0.027], Fig. 4B), suggestive of individual differences with a group-level mean close to 0. We performed several model validation checks to verify that the winning model M12 was able to capture key qualitative features of the empirical data (posterior predictive checks, Fig. 4C), could identify data-generating parameters reliably (parameter recovery, Fig. 4D), and could be distinguished from other models (model recovery, Fig. 4E, see also Supplementary Material S07).

**Figure 4.**
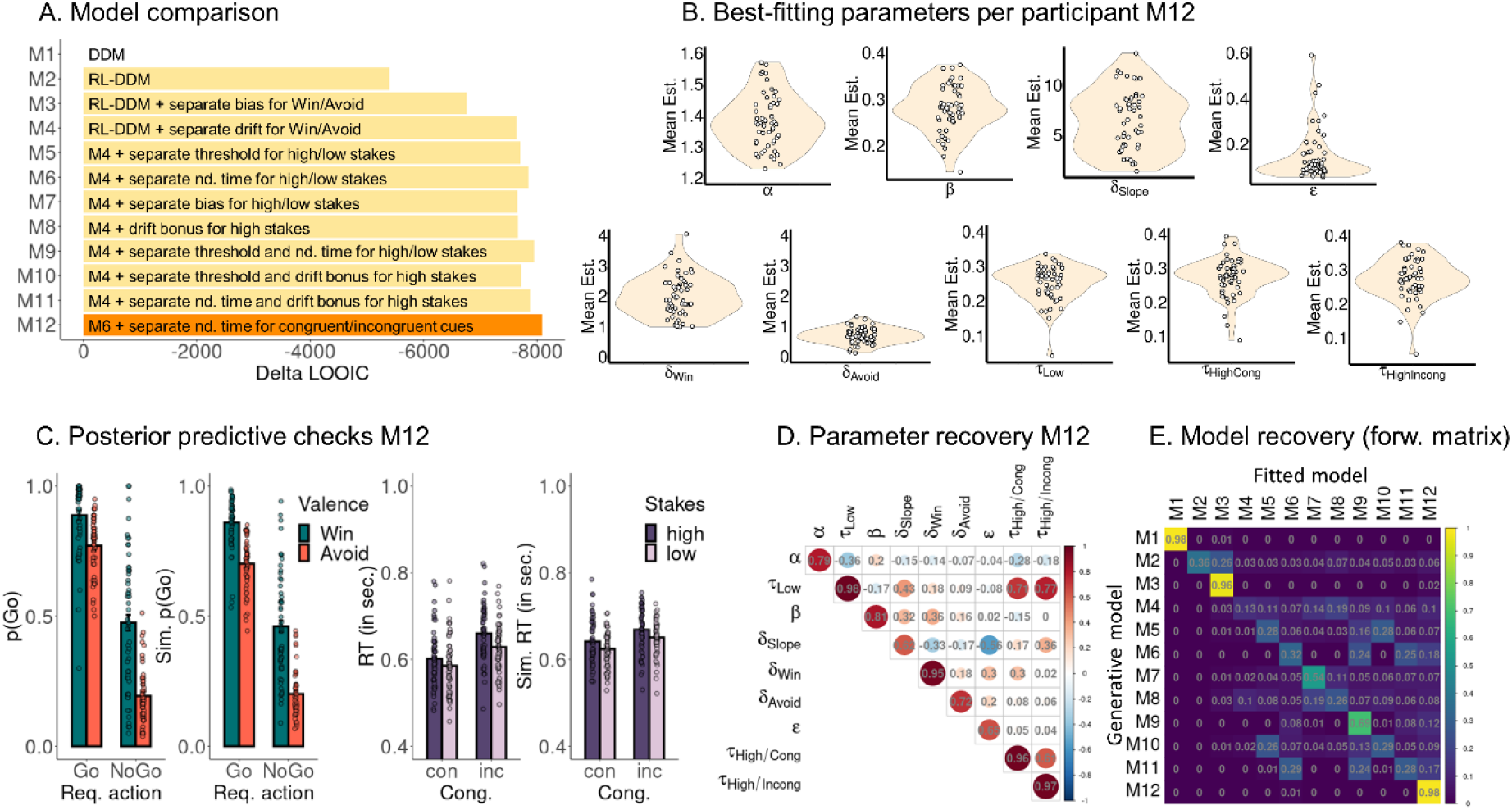
Reinforcement-learning drift-diffusion models. **A.** Model comparison. LOO-IC favors model M12, implementing separate drift rate intercepts for Win and Avoid cues and separate non-decision times for low stakes, congruent cues under high stakes, and incongruent cues under high stakes. **B.** Densities of best fitting parameters for model M12 per participant. Drift rate intercepts for Win cues are consistently higher than drift rate intercepts for Avoid cues. Note that, although the winning model implements separate non-decision times for high/low stakes and congruent/incongruent cues, the parameter values for these different conditions are not significantly different from each other. **C.** Posterior predictive checks for the winning model M12. Left panel: Simulated proportion of Go responses per required action and cue valence averaged over simulations and participants. The winning model M12 reproduces Pavlovian biases in responses and RTs (see Supplementary Material S07). Right panel: Simulated RTs per cue congruency per stakes level averaged over simulations and participants. The winning model M12 reproduces the overall slowing under high stakes as well as differences in slowing between congruent and incongruent cues, but somewhat underestimates this difference compared to the empirical data (see Supplementary Material S07 for further plots). For further plots, see Supplementary Material S07. **D.** Parameter recovery for the winning model M12. Correlations between generative parameters used for simulating 1,000 data sets based on M12 and parameters obtained when fitting M12 to simulated data. All correlations between generative and fitted parameters (on-diagonal correlations) are significantly above chance (*Mr* = 0.83, *SDr* = 0.14, range 0.62–0.98; 95^th^ percentile of permutation null distribution: *r* = 0.08; see Supplementary Material S07 for scatter plots of on-diagonal correlations). Besides correlations between generative parameters with their corresponding fitted parameters, there were two notable cases of off-diagonal correlations: first, the different non-decision times (under low stakes, under high stakes for congruent cues, and under high stakes for incongruent cues) were correlated (*r* = 0.71 and *r* = 0.77), reflecting an overall tendency towards faster/slower responses that is naturally shared across all three parameters. Second, learning rates and drift rate slopes were negatively correlated across parameter settings (*r* = -0.56), which mimics the frequently observed trade-off between learning rate and inverse temperature parameters in more classic reinforcement learning models of choices (Ballard and McClure 2019). In RL-DDMs, the drift rate slope is multiplied with the Q-value difference, such that steeper slopes lead to more deterministic choices and shallower slopes lead to more stochastic choices, similar to an inverse temperature parameter. **E.** Model recovery for model M1-M12. The forward confusion matrix displays the conditional probabilities that model Y is identified as the best fitting model (columns) if model X (rows) is the underlying generative model used to simulate a given data set. On-diagonal probabilities indicate the probability of reidentifying the generative model. All on-diagonal probabilities are significantly above chance (*Mp* = 0.31, *SDp* = 0.32, range 0.13–0.98; 95^th^ percentile of permutation null distribution: *p* = 0.10). Model recovery was particularly high for the winning model M12, which was the best fitting model for 98% of data sets for which it was the generative model. Recovery for the other models was not quite as high, though still significantly above chance for all models. Hence, while our model selection procedure will likely identify M12 if it is the data generating process, it might struggle in other scenarios in which other DDM parameters are affected by experimental manipulations. For the inverse confusion matrix and matrices on subsets of models, see Supplementary Material S07.

In sum, model comparison results were in line with the regression results, yielding a selective effect of stakes in prolonging the non-decision time, and separately so for incongruent and congruent cues. Stakes did not affect the threshold and/or the drift rate as typically observed in a speed-accuracy trade-off. Hence, we conclude that stakes do not shift the speed-accuracy trade-off, but rather lead to response slowing independent of response selection.

## Discussion

In this pre-registered experiment, we found evidence that increasing stake magnitude slowed down responses in a Motivational Go/NoGo Learning Task, especially for incongruent cue conditions, without affecting whether participants responded or not. In line with previous literature, participants exhibited a Pavlovian bias in both responses and RTs (Algermissen et al. 2022; Swart et al. 2017), with more and faster Go responses to Win than Avoid cues. On trials with high stakes (i.e., larger rewards or punishments at stake), they slowed down, particularly for the two incongruent conditions Go-to-Avoid and NoGo-to-Win. This response slowing was best described by high stakes prolonging the non-decision time in a drift-diffusion model framework, particularly so for incongruent trials. This finding is inconsistent with both hypotheses put forward in the introduction, i.e., high stakes strengthening Pavlovian biases or high stakes motivating cognitive control to suppress them on incongruent trials. In sum, higher stakes slow down response selection, but neither strengthen nor weaken Pavlovian biases in responses. We propose two possible explanations for this (somewhat surprising) result: response slowing under high stakes might reflect (flexibly recruited) cognitive control, which is however ineffectively used; or it might reflect (automatic/reflexive) positive condition suppression, i.e., the suppression of goal-directed behavior by large imminent rewards as previously observed in animal studies.

On trials with high stakes, participants took longer to make a Go response, but did not exhibit any altered tendency for Go/NoGo responses, i.e. no reduction or enhancement of Pavlovian biases. Apart from the null effect on responses, RTs slowed down under high stakes, an effect that was highly consistent across participants (Fig. 3E, F). These two findings are incompatible with the first hypothesis posited, i.e., high stakes strengthening Pavlovian biases. Slowing (instead of speeding) of responses under high rewards might appear quite surprising given a large body of literature showing higher stakes speed up responses (Fontanesi et al. 2019; Luo et al. 2009; Pirrone et al. 2018; Smith and Krajbich 2018) and some evidence for larger PIT effects for high compared to low value cues (Algermissen & den Ouden, 2023; Schad et al., 2020). Notably, response slowing occurred for both appetite and aversive cues, suggesting that the effect is independent of cue valence and orthogonal to the Pavlovian biases. Note that 50% of trials were high stake trials, arguing against the possibility of surprise (i.e., oddball effects) driving the response slowing. High and low stake trials were visually very distinct, arguing against differences in processing demands between both trial types. In sum, the size of Pavlovian biases in the Motivational Go/NoGo Task appears to be unaffected by stake magnitude, which instead induced a response slowing orthogonal to the biases.

Response slowing under high stakes might be partly compatible with the second hypothesis (EVC), i.e., high stakes increasing cognitive control in order to suppress biases, given that heightened cognitive control recruitment is often inferred from/accompanied by prolonged reaction times (Frank 2006; Shenhav et al. 2013; Wessel and Aron 2017). Specifically, in line with our preregistered hypothesis that high stakes increase cognitive control recruitment, response slowing was stronger on motivationally incongruent trials on which Pavlovian biases had to be suppressed in order to execute the correct response. This effect suggests that participants did distinguish the different cue conditions with respect to whether they could benefit from increased cognitive control recruitment and prolonged deliberation times (i.e., situations in which control could in theory change the emitted response) or not.

One possibility is that, in fact, both hypotheses are true and high stakes strengthen biases, but also lead to heightened control recruitment countering the biases. Both effects would cancel out in the responses (Go/NoGo) and only remain visible in prolonged reaction times. In this scenario, extra control recruitment might have prevented a further escalation of the biases under high stakes. However, if conflict is detected and additional control recruited on the appropriate subset of (incongruent) trials, it is unclear why not even more control is recruited to reduce biases to zero. Future studies using independent measures of incentive motivation and cognitive control recruitment, such as fMRI, EEG, or pupillometry (Algermissen et al. 2022; Algermissen and den Ouden 2024) could test this interpretation. In absence of such evidence, the most parsimonious conclusion is that, in this data set, high stakes led to increased cognitive control recruitment, but control was ineffective at improving response selection, with the size of Pavlovian biases (in terms of the proportion of Go responses to Win and Avoid cues) unaltered under high stakes.

Cognitive control recruitment is usually regarded as a deliberate, strategic process that only occurs when a cost-benefit tradeoff favors additional control (Shenhav et al. 2013). However, some studies have reported response slowing that seemingly reflects cognitive control, but occurs in situations in which it does not help task performance. Such findings suggest that response slowing can occur in an automatic, reflexive manner with limited sensitivity to the task context. One example is the phenomenon of post-error slowing, i.e., participants showing slower RTs on trials following incorrect responses than trials following correct responses, which is typically interpreted as a proactive cognitive control process to prevent future errors (Wessel 2018). However, in very difficult tasks with more incorrect than correct responses, this phenomenon reverses, with slower responses after correct than incorrect trials, suggesting that slowing occurs reflexively after rare events even if it is not conducive to future performance (Notebaert et al. 2009; Núňez Castellar et al. 2010). Similarly, participants have been found to overexert control for a selection of stimuli in a Stroop task if these stimuli had previously been paired with high rewards, even if this reward had subsequently been removed (Bustamante et al. 2021). Together, these results suggest that response slowing (putatively reflecting cognitive control recruitment) is sometimes shown in a non-strategic, reflexive manner in situations in which it does not help task performance. We speculate that, in our task, similarly, higher stakes lead to a reflexive slowing of responses, even if this slowing does not help overcome Pavlovian biases.

An alternative explanation for response slowing under high stakes might be the phenomenon of “choking under pressure”, i.e., the fear of failure in high-stakes situations inducing rumination and thus decreasing performance (Beilock and Carr 2001, 2005; Chib et al. 2012, 2014; Dunne et al. 2019), an option we had considered in our pre-registration. Choking under pressure predicts a pattern opposite to the second hypothesis (EVC), with high stakes undermining cognitive control recruitment and leading to lower performance in incongruent conditions. While the observed slowing of RTs could be interpreted as a kind of “choking under pressure”, we did not observe corresponding performance decrements. Hence, this finding does not fall under the phenomenon of “choking under pressure” as investigated in previous literature. In sum, these results are most compatible with the idea of high stakes leading to increased cognitive control recruitment, though without any consequences for response selection and accuracy.

Past computational models have proposed mechanisms of how decision accuracy—which is particularly warranted in high stakes situations—can be prioritized over speed by increasing decision thresholds in an evidence accumulation framework (Bogacz et al. 2006). Such increased decision bounds have been typically investigated in situations in which choice options are close in value and thus eliciting cognitive conflict. Neuro-computational models suggest that such conflict is detected by the anterior cingulate cortex and presupplementary motor area, which—via the hyperdirect pathway involving the subthalamic nucleus—project to the globus pallidus and increase decision thresholds in the basal ganglia action selection circuits, leading to a higher requirement for positive evidence to elicit a response (Cavanagh et al. 2011; Forstmann et al. 2008; Frank 2006; Frank et al. 2015; Wiecki and Frank 2013). This decision threshold adjustment will lead to a higher proportion of correct, but overall slower responses. It is plausible that the same mechanism could lead to response caution in the context of high-value cues. In fact, a series of recent studies found that cues indicating an upcoming choice between high-value options (but not the presence of high-value options per se) slowed down of RTs, which was best captured by a heightened decision threshold (Shevlin et al. 2022). However, in contrast, the data of the present study were best explained by a model embodying prolonged non-decision times rather than heightened response thresholds. It is thus unclear whether the same computational and neural mechanisms proposed for implementing speed-accuracy tradeoffs are also responsible for the response slowing observed in this data. Future studies using neuroimaging of cortical and subcortical activity (Algermissen et al. 2022) and instructions to prioritize speed or accuracy during the task (Forstmann et al. 2008) while simultaneously manipulating stakes could shed light on shared vs. separate neural mechanisms.

Another possible interpretation of our findings is that the response slowing under large stake magnitudes is an instance of positive conditioned suppression as previously reported in rodents (Azrin and Hake 1969; Van Dyne 1971; Timberlake et al. 1982). In positive conditioned suppression, cues signaling the imminent receipt of a reward globally suppress behavior. Specifically, a cue announcing an imminent reward suppresses reflexive exploratory behaviors that would move the animal away from a food site, and instead invigorates and prolongs engagement with the site of reward delivery until the reward is obtained (Marshall et al. 2020). This phenomenon might be adaptive in that it prevents agents from becoming distracted by other reward opportunities and forgetting to collect the reward they previously worked for (Konorski 1967; Timberlake et al. 1982). However, for sufficiently large rewards, this suppression might extend backwards in time such that it even affects the instrumental response required to obtain the reward (i.e., a lever press) (Marshall and Ostlund 2018; Marshall et al. 2023). Hence, positive conditioned suppression seems to arise from a conflict between preparatory behaviors used to obtain an outcome and consummatory behaviors needed to collect it, and might only arise for relatively large outcomes. In line with this account, a recent animal study found small rewards to invigorate responding in line with classical PIT findings (Marshall et al. 2023). However, large rewards suppressed instrumental lever pressing and diminished PIT effects, suggestive of positive conditioned suppression interfering with PIT in a way similar to our findings.

Interpreting response slowing as positive conditioned suppression implies that it should occur automatically and reflexively, with no sensitivity for the particular task context. However, the prolongation of RTs in the present data was particularly strong for motivationally incongruent cues, which argues against a purely automatic, “reflexive” nature of the observed effect, and instead in favor of an adaptive effect that is (at least partially) sensitive to task requirements. It is possible that both (automatic) positive conditioned suppression and (voluntary) heightened cognitive control recruitment triggered by motivational conflict are present in tandem, or that positive conditioned suppression is (partially) a consequence of cognitive control recruitment. While both conditioned suppression and cognitive control are sensitive to stakes, conditioned suppression should be uniquely sensitive to the delay between the response and outcome delivery because it reflects interference between preparatory and consummatory processes. Hence, future studies could try to disentangle both interpretations of our data by varying the delay between response execution and outcome delivery in a predictable fashion. Stronger slowing for shorter delays would uniquely support the interpretation of positive conditioned suppression (Marshall et al. 2023; Marshall and Ostlund 2018; Meltzer and Hamm 1978; Miczek and Grossman 1971).

Furthermore, conditioned suppression has yet not been studied in the context of avoiding aversive outcomes. Slowing induced by conditioned suppression will look highly similar to slowing induced by the Pavlovian bias itself. In our data, effects of action-valence congruency (i.e. Pavlovian bias) and stake magnitude on RTs were additive, suggesting independent mechanisms. Future research might try to disentangle these two effects further by using an “escape” context in which participants must select actions to terminate an ongoing aversive stimulus (e.g. loud noise). When terminating an ongoing aversive stimulus, participants show a tendency towards Go (“aversive invigoration”) while, when aiming to prevent a future punishment, they show a tendency towards NoGo (“aversive inhibition”) (Millner et al. 2017). In an escape scenario, if high stakes invigorated an existing Pavlovian bias, we would expect more Go responses, while if high stakes led to positive conditioned suppression, we would expect fewer Go responses. Such an experiment could reveal if high stakes merely potentiate an existing bias or lead to positive conditioned suppression as an independent, additive phenomenon.

The ability to inhibit impulsive behaviors has been proposed to be serotonergic in nature. Serotonin agonists slow down behavior (Soubrié 1986), while serotonin depletion speeds up responses and can lead to disinhibition and impulsivity (Boureau and Dayan 2011; Dalley and Roiser 2012; Bari and Robbins 2013). Inhibition is usually observed in the aversive domain, preventing responses to aversive stimuli, which become disinhibited under serotonin depletion (Crockett et al. 2009, 2012; Geurts et al. 2013b). However, serotonergic neurons also respond to rewards (Kranz et al. 2010; Luo et al. 2015; Liu et al. 2020). An influential theory suggests that serotonin aids the suppression of automatic reflexes in service of goal-directed behaviors via its effects on the periaqueductal gray (Deakin and Graeff 1991; Graeff 2002, 2004). This proposed function is remarkably similar to positive conditioned suppression and could serve both punishment prevention and reward pursuit. High stakes might activate serotoninergic circuits and prevent reflexive escape responses to aversive stimuli as well as reflexive approach responses to minor appetitive stimuli. Indeed, we have previously observed that participants slow down when stakes are high, but that this slowing is abolished under serotonin depletion (den Ouden et al. 2015). Animal studies have found that the activation of serotonergic neurons facilitates waiting for rewards (Miyazaki et al. 2011, 2020, 2014) and persistence in foraging (Lottem et al. 2018). In sum, high stakes might activate serotoninergic circuits and in this way suppress reflexive responses in service of goal-directed behavior, which leads to a general slowing in the context of our task. Future research should explicitly test the putatively serotonergic nature of high stakes-induced response slowing in the Motivational Go/NoGo Task using pharmacological manipulation and/or recordings from serotonergic neural circuits.

A limitation of the current study is that high stakes were explicitly signaled via a red circle around the task cue. In this way, the task mimicked situations in which simple visual features tell apart high stakes from low stakes situations, e.g. a lion from a spider. However, it does not mimic situations in which high value must be inferred indirectly from past experiences or by combining multiple features, e.g., when selecting the best bargain among a range of houses or cars. It is possible that response slowing only occurs when high stakes situations are explicitly cued (Shevlin et al. 2022), whereas high stakes invigorate Pavlovian biases when stakes have to be inferred from a combination of visual features (Knutson et al. 2005), are relevant only for a secondary task (Algermissen and den Ouden 2023) or completely task-irrelevant such as in many versions of the PIT task (Geurts et al. 2013a, 2013b; Schad et al. 2020). This is an important consideration for task designs that might explain the mixed literature on stakes effects in PIT tasks. Finally, the presented finding mimics cases where “high stakes” describes the entire situation rather than a single option (Shevlin et al. 2022), but is unlike cases where only a single option is more valuable and dominates all other options.

Another limitation might be that stakes were not varied in a continuous fashion, but categorically as two discrete levels. Again, it might be plausible that agents represent situations (e.g. trials) as overall “high stakes” or not, irrespective of the particular value of single options (Shevlin et al. 2022). Varying the stakes magnitude in a continuous fashion would increase processing demands and thus already slow down responses due to perceptual (irrespective of additional decision) difficulty. Furthermore, participants might subjectively recode stakes levels relative to the mean stake level, representing low rewards as disappointing and thus akin to punishments, while perceiving low punishments as a relief and thus akin to rewards (Klein et al. 2017; Palminteri et al. 2015). These considerations support the ecological validity of dichotomizing stakes into high and low levels. However, it remains to be empirically tested whether continuous stakes levels lead to similar or different effects.

In sum, while possibilities to gain rewards/avoid punishments induce Pavlovian biases, increasing the stakes of these prospects does not alter the strength of biases. However, high stakes motivate humans to slow down their responses. One interpretation is that this slowing is adaptive in allowing time for conflict detection and cognitive control recruitment in case motivational biases have to be suppressed. However, the slowing is not associated with changes in response selection, i.e., also not with the degree to which participants suppress their Pavlovian biases when these are unhelpful, suggesting that humans do not use this additional time effectively. An alternative interpretation is that prolonged reaction times reflect positive conditioned suppression, i.e. interference between goal-directed and consummatory behaviors triggered by large stakes as previously observed in rodents. Taken together, this study suggests that high stakes might have a similar effect in both humans and rodents in the context of Pavlovian/instrumental interactions on action selection.

## Methods

### Participants and Exclusion Criteria

Fifty-five human participants (*M*age = 22.31, *SD*age = 2.21, range 18–29; 42 women, 13 men; 47 right-handed, 8 left-handed) participated in an experiment of about 45 minutes. The study design, hypotheses, and analysis plan were pre-registered on OSF under https://osf.io/ue397. Individuals who were 18–30 years old, spoke and understood English, and did not suffer from colorblindness were recruited via the SONA Radboud Research Participation System of Radboud University. Their data were excluded from all analyses for two (pre-registered) reasons: (a) guessing the hypotheses of the experiment on the first question of the debriefing, which was not the case for any participant; (b) performance not significantly above chance (tested by using required action to predict performed action with a logistic regression; only participants with *p* < .05 were included), which was the case for one participant. All the results presented in the main text are thus based on a final sample of *N* = 54. See the Supplementary Material S03 for results based on all 55 participants, which led to identical conclusions. This research was approved by the local ethics committee of the Faculty of Social Sciences at Radboud University (proposal no. ECSW-2018-171) in accordance with the Declaration of Helsinki.

The sample size was not based on a power analysis, but on lab availability for this project (three weeks). This study was conducted as part of final year thesis projects, which received special lab access in this period. The final sample size of *N* = 54 was larger than previous studies investigating Pavlovian biases with the same task (Algermissen et al. 2022; Swart et al. 2018) and more than twice as large as comparable studies investigating the effect of incentives on cognitive control recruitment (Chiew and Braver 2016; Krebs et al. 2010). A post-hoc sensitivity power analysis yielded that, given 54 participants providing 320 trials, thus 17,280 trials in total, assuming an intra-cluster coefficient of 0.043 for responses and 0.094 for RTs (estimated from the data), the effective sample size was *n* = 5,281 for responses and *n* = 2,877 for RTs, which allowed us to detect effects of β > .039 (standardized regression coefficient) for responses and β > .052 for RTs with 80% power (Aarts et al. 2014).

### Procedure

Participants completed a single experimental session that lasted about 45 minutes. After providing informed consent, participants received computerized instructions and performed four practice trials for each of the four task conditions. Afterwards, they completed 320 trials of the Motivational Go/NoGo Task. After the task, participants performed the V5-D MESA Digit Span Test measuring forward and backward digit span (Fitzpatrick et al. 2015) and filled in the non-planning subscale of the Barratt Impulsiveness Scale (Patton et al. 1995) and the neuroticism sub-scale of the Big Five Aspects Scales (DeYoung et al. 2007). These measures were part of final year thesis projects and not of focal interest (see the pre-registration); results are reported in Supplementary Material S05. Finally, participants went through a funnel debriefing asking them about their guesses of the hypothesis of the study, whether they used specific strategies to perform the task, whether they found the task more or less difficult to perform on high stakes trials, and if so, whether they had an explanation of why this was the case. At the end, they received course credit for participation as well as a small extra candy reward when they scored more than 960 points (equivalent to 67% accuracy across trials, equivalence unknown to participants), which was announced in the instructions.

### Task

Participants completed 320 trials (80 per condition; 40 each with high and low stakes respectively) of the Motivational Go/NoGo learning task. Each trial started with one of four abstract geometric cues presented for 1,300 ms (Fig. 1A). The assignment of cues to task conditions was counterbalanced across participants. Participants needed to learn from trial-and-error about the cue valence, i.e., whether the cue was a Win cue (point gain for correct responses; no change in point score for incorrect responses) or an Avoid cue (no change in point score for correct responses; point loss for incorrect responses), and the required action, i.e., whether the correct response was Go (a key press of the space bar) or NoGo (no action; Fig. 1B). Participants could perform Go responses while the cue was on the screen. In 50% of trials, the cue was surrounded by a dark red circle (RGB [255, 0, 0]), signaling the chance to win or avoid losing 50 points (high stakes condition). On all other trials, 10 points could be won or lost (low stakes condition). After a variable inter-stimulus interval of 500–900 ms (uniform distribution in steps of 100 ms), numerical feedback was presented for 700 ms (+10/+50 in green font for point wins, -10/-50 in red font for point losses; 000 in grey font for no change in point score). Feedback was probabilistic such that correct responses were followed by favorable outcomes (point win for Win cues, no change for Avoid cues) on only 80% of trials, while on the other 20% of trials, participants received unfavorable outcomes (no change for Win cues, point loss for Avoid cues; Fig. 1C). These probabilities were reversed for incorrect responses. Probabilistic feedback was used to make learning more difficult and induce a slower learning curve. Trials ended with a variable inter-trial interval of 1,300–1,700 ms (uniform distribution in steps of 100 ms).

The task was administered in four blocks of 80 trials each. Each block featured a distinct set of four cues for which participants had to learn the correct response. Probabilistic feedback and renewal of the cue set were used to slow down learning, given previous findings that biases disappear when accuracy approaches 100% (Swart et al. 2017).

## Data Analysis

### Data Preprocessing

(Trials with) RTs faster than 300 ms (regardless of whether the response was correct or not) were excluded from all analyses as those were assumed to be too fast to reflect processing of the cue. This was the case for 103 out of 17,600 trials (0.6% of trials; per participant: *M* = 1.91, *SD* = 5.89, range 0–41). We did not exclude error trials because these are needed to assess both learning and the presence of Pavlovian biases. See Supplementary Material S02 for results using all reaction times from all trials.

### Mixed-effects Regression Models

We tested hypotheses using mixed-effects linear regression (function lmer) and logistic regression (function glmer) as implemented in the package lme4 in R (Bates et al. 2015). We used generalized linear models with a binomial link function (i.e., logistic regression) for binary dependent variables such as accuracy (correct vs. incorrect) and response (Go vs. NoGo), and linear models for continuous variables such as RTs. We used zero-sum coding for categorical independent variables. All continuous dependent and independent variables were standardized such that regression weights can be interpreted as standardized regression coefficients. All regression models contained a fixed intercept. We added all possible random intercepts, slopes, and correlations to achieve a maximal random effects structure (Barr et al. 2013). *P*-values were computed using likelihood ratio tests with the package afex (Singmann et al. 2018). We considered *p*-values smaller than α = 0.05 as statistically significant.

### Evidence for absence of an effect

We plot the condition means for each participant and provide confidence intervals for every effect. Every possible point estimate of an effect that would fall outside the estimated confidence interval can be rejected at a level of α = 0.05.

### Computational modeling of responses and reaction times

#### Combining reinforcement learning with a drift-diffusion choice rule

A class of computational models that allows to jointly model both responses and reaction times are so called “evidence accumulation” or “sequential sampling” models such as the drift-diffusion model (DDM) (Ratcliff 1978). These models formalize a decision process in which evidence for two (or more) response options is accumulated until a fixed threshold, and a response is elicited upon reaching this threshold. The process is captured through four parameters (Wabersich and Vandekerckhove 2014): the drift rate δ, reflecting the speed with which evidence is accumulated; the decision threshold α, describing the distance of the threshold from the starting point; the starting point bias β, reflecting if the accumulation process starts in the middle between both bounds (β = 0.5) or closer to one of the boundaries, reflecting an overall response bias; and the non-decision time τ; capturing the duration of all perceptual or motor processes that contribute to RT, but are not part of the decision process itself.

Typically, DDMs aim to explain choices when response requirements given a certain visual input are clear to the participant. However, in the current study, participants learn the correct response for each cue over time, leading to progressively faster and more accurate responses. Recent advances in computational modeling propose that it is possible to combine drift-diffusion models with a reinforcement learning (RL) process, yielding a reinforcement-learning drift-diffusion model (RL- DDM) (Fontanesi et al. 2019; Pedersen et al. 2017; Miletić et al. 2020). We employed a simple Rescorla-Wagner model which uses outcomes r (+1 for rewards, 0 for neutral outcomes, -1 for punishments) to compute prediction errors r – Q, which we then used to update the action value Q for the chosen action a towards cue s:

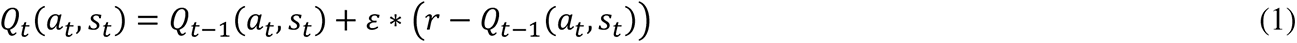

Here, the difference in Q-values between choice options (QGo – QNoGo) serves as the input to the drift rate. This difference is initially zero, but grows with learning (positive difference if “Go” leads to more rewards, and negative difference if “NoGo” leads to more rewards). This Q-value difference is then multiplied with a constant drift rate parameter. At the beginning of the learning process, the resulting low drift rates lead to more stochastic choices and slow RTs, but, as the Q-value difference grows, higher drift rates result in more deterministic choices and faster RTs. The learning process requires an additional free parameter, i.e., the learning rate parameter ε, which determines the impact of the prediction error on belief updating. The drift rate parameter acts akin to the inverse temperature parameter used in the softmax choice rule, with higher drift rates leading to more deterministic choices.

One peculiarity of the Motivational Go-NoGo Task is the NoGo response option, which by definition does not yield RTs. Variants of the DDM allow for such responses by integrating over the latent RT distribution of the implicit NoGo decision boundary (Gomez et al. 2007; Ratcliff et al. 2018), for which an approximation exists (Blurton et al. 2012). This implementation has previously been used to model another variant of motivational Go/NoGo task (Millner et al. 2017) and is implemented in the HDDM toolbox (Wiecki et al. 2013).

Note that RL-DDMs were not mentioned in the pre-registration, which only mentioned reinforcement learning models to-be fitted to participants’ choices. In light of the results from the regression analyses, incorporating RTs into the model and testing alternative mechanisms by which stakes could influence the choice process seemed warranted.

#### Model space

We fit a series of increasingly complex models. We first tested whether an RL- DDM fit the data better than a standard DDM; then tested the computational implementation of the Pavlovian bias, and lastly tested the effect of stakes on model parameters. Model **M1** (parameters α, τ, β, δINT) just featured the DDM model with a constant drift rate parameter, but no learning, assuming that participants have a constant propensity to make a Go response for any trial, irrespective of the presented cue. **M2** (parameters α, τ, β, δINT, δSLOPE, ε) added a reinforcement learning process, updating Q-values for Go and NoGo for each cue with the observed feedback, multiplying the Q-value difference (QGo – QNoGo) with the drift rate parameter δSLOPE and finally adding it to the drift-rate intercept δINT to obtain the net drift rate. Including a drift-rate intercept δINT, i.e., an overall tendency towards making a Go/NoGo response even when the Q-value difference was zero, which is similar to an overall Go bias parameter, yielded considerably better fit than models without such an intercept. If people learned the task, model M2 should fit their data better than M1. Next, M3 and M4 comprised different implementations of the Pavlovian bias, either assuming separate starting point biases (**M3**; parameters α, τ, βWIN, βAVOID, δINT, δSLOPE, ε) or alternatively separate drift rate intercepts (**M4**; parameters α, τ, β, δWIN, δAVOID, δSLOPE, ε) for Win and Avoid cues, two plausible implementations considered in previous literature (Millner et al. 2017). Next, models M5-M8 (parameters α, τ, β, δWIN, δAVOID, δSLOPE, ε, one additional parameter π for high stakes) extended M4 and tested possible effects of the stakes on a single parameter, implementing effect of the stakes on the threshold (**M5**), the non-decision time (**M6**), the bias (**M7**) and the drift rate intercept (**M8**). As a control, models M9-M11 (parameters α, τ, β, δWIN, δAVOID, δSLOPE, ε, two additional parameters π and θ for high stakes) tested effects of stakes on two parameters (only combinations that could potentially give rise to response slowing), namely on both the threshold and the non-decision time (**M9**), the threshold and the drift rate (**M10**; i.e. the two parameters typically modulated by speed-accuracy trade-offs), and the non-decision time and drift rate (**M11**).

Finally, given the results from model comparison of these earlier models, **M12** tested whether the effect of stakes of non-decision time was different for congruent and incongruent cues.

#### Priors, transformations, parameterization, and starting values

We fitted models in a hierarchical Bayesian fashion, modeling group-level parameters (means and standard deviations) that served as priors for the subject-level parameters using the probabilistic programming language Stan (Carpenter et al. 2017) in R (rstan). Stan implements a Hamiltonian Monte-Carlo (HMC) Markov-chain algorithm with a No-U-Turn sampler (NUTS). We used the following group-level hyperpriors: Mδ ∼ *N*(5, 2), Mα ∼ *N*(0, 1), Mβ ∼ *N*(0, 1), Mτ ∼ *N*(0, 1), Mε ∼ *N*(0, 1), Mπ ∼ *N*(0, 1), Mϑ ∼ *N*(0, 1), and for all SDs: SD ∼ *N*(0, 1). The parameters δ, α, τ were constrained to be positive by using the y = log(1 + exp(x)) transformation, which is y = 0 for negative numbers, smoothly asymptotes 0 for small positive numbers, and is roughly y = x for large positive numbers. The parameters β and ε were constrained to be in the range [0, 1] by using a softmax transformation y = exp(x) / (1 + exp(x)). In line with previous DDM implementations in Stan (Fontanesi et al. 2019; Kraemer et al. 2021), we used a non-centered parameterization in which individual-subject parameters are modeled with a standard normal prior *N*(0, 1) that is first multiplied with the group-level standard deviation and then added to the group-level mean parameter. Furthermore, again in line with previous DDM implementations in Stan (Fontanesi et al. 2019; Kraemer et al. 2021), we set the following starting values: *M*α = -0.18, *M*τ = -10, Mβ ∼ *N*(0.5, 0.1), MδINT ∼ *N*(0, 1), MδSLOPE∼ *N*(0, 1), Mπ ∼ *N*(0, 0.1), Mθ ∼ *N*(0, 0.1), all group-level *SDs* = 0.001, all subject level parameters as ∼ *N*(0, 1). For models with an effect of stakes on the non-decision-time (M6, M9, M11), τ (low stakes) had to be initialized to be considerably smaller than π (high stakes), which was accomplished by Mτ ∼ *N*(0, 1e-6) and SDτ = 1e-6.

#### Model fitting and convergence checks

For each model, we used four chains with 10,000 iterations each (5,000 as warm-up), yielding a total of 20,000 samples contributing to the posteriors. We checked that Rhats for all parameters were below 1.01, effective sample sizes for all parameters were at least 400, that chains were stationary and well-mixing (using trace plots), that the Bayesian fraction of missing information (BFMI) for each chain was above 0.2, and that (if possible) no divergent transitions occurred (Baribault and Collins 2023). To minimize the occurrence of divergent transitions, we increased the target average proposal acceptance probability (adapt_delta) to 0.99. We visually inspected that posterior densities were unimodal and no strong trade-offs between parameters across samples occurred.

#### Model comparison

For model comparison, we used the LOO-IC (efficient approximate leave-one-out cross-validation information criterion) based on Pareto-smoothed importance sampling (PSIS) (Vehtari et al. 2017). For completeness, we also report the WAIC (widely applicable information criterion) in Supplementary Material S07, but give priority to the LOO-IC, which is more robust to weak priors or influential observations (Vehtari et al. 2017). Both WAIC and LOO-IC behave like the negative log-likelihood, with lower numbers indicating better model fit.

#### Posterior predictive checks

For the winning model M12, we randomly drew 1,000 samples from the posteriors of each participants’ subject-level parameters, simulated a data set for each participant for each of these 1,000 parameter settings, and computed the mean simulated p(Go), p(Correct), and RT for each participant for each trial across parameter settings. We then plotted the mean simulated p(Go), p(Correct), and RT as a function of relevant task conditions to verify that the model could reproduce key qualitative patterns from the empirical data (Palminteri et al. 2017).

#### Parameter recovery

For the winning model M12, we fitted a multivariate normal distribution to the mean subject-level parameters across participants and sampled 1,000 new parameter settings from this distribution. We simulated a data set for each parameter setting and fitted model M12 to the simulated data. We then correlated the “ground-truth” generative parameters used to simulate each data set to the fitted parameters obtained when fitting M12 to it. To evaluate whether correlations were significantly higher than expectable by chance, we computed a permutation null distribution of the on-diagonal correlations. For this purpose, over 1,000 iterations, we randomly permuted the assignment of fitted parameter values to data sets, correlated generative and fitted parameter values, and saved the on-diagonal correlations. We tested empirical correlations against the 95^th^ percentile of this permutation null distribution.

#### Model recovery

For each of the 12 models, we fitted a multivariate normal distribution to the mean subject-level parameters across participants and sampled 1,000 new parameter settings from it (with the constraints that learning rates were required to be > 0.05 and parameter differences sampled from the upper 50% of the parameter distribution to keep models distinguishable). We simulated a new data set for each parameter setting, resulting in total in 12,000 data sets. We fitted each of the 12 models to each data set, resulting in 144,000 model fits. For each data set, we identified the model with the lowest LOO-IC. We counted how often each fitted model Y emerged as the winning model for the data sets of each generative model X, computing the forward confusion matrix containing conditional probabilities p(best fitting model = Y | generative model = X) for each combination of generative model X and fitted model Y (Wilson and Collins 2019). We also computed the inverse confusion matrix containing p(generative model = X | best-fitting model = Y; see Supplementary Material S07). To evaluate whether these probabilities were significantly higher than expectable by chance, we computed a permutation null distribution of the on-diagonal probabilities. For this purpose, over 1,000 iterations, we randomly permuted the LOO-IC values of all fitted models for a given data set, counted how often each fitted model emerged as the winning model for the data sets of each generative model, and extracted the on-diagonal probabilities. We tested empirical probabilities against the 95^th^ percentile of this null distribution.

### Transparency and openness

We report how we determined our sample size, all data exclusions, all manipulations, and all measures in the study. All data, analysis code, and research materials will be shared upon publication. The study design, hypotheses, and analysis plan were pre-registered on OSF under https://osf.io/ue397. Data were analyzed using R, version 4.1.3 (R Core Team, 2022). Models were fitted with the package lme4, version 1.1.31 (Bates et al., 2015). Plots were generated with ggplot, version 3.4.2 (Wickham, 2016).

## Conflict of interest

We have no known conflict of interest to disclose.

## Funding

J. Algermissen was funded by a PhD position from the Donders Centre of Cognition, Faculty of Social Sciences, Radboud University, the Netherlands. Hanneke E.M. den Ouden was supported by a Netherlands Organization for Scientific Research (NWO) VIDI grant 452-17-016.

## Acknowledgements

We thank Theresa Altmeyer, Max Molhuizen, Eva Saragosa, and Madeleine Stahlhacke for assistance with data collection.

## Supplemental Material S01: Overview results mixed-effects regression models

Here, we report an overview over all major statistical results reported in the main text and the supplementary material. For details on how mixed-effects regression were performed, see the Methods section of the main text.

**Table S01.** *Overview of the results from all mixed-effects regression models reported the main text of the manuscript*. Featuring data from *N* = 54 participants, trial with RTs < 0.300 sec. are excluded from RT analyses

| Model ID | DV | IV | b | SE | $\chi^2(1)$ | p |
| --- | --- | --- | --- | --- | --- | --- |
| 1 | Response | Required action | 1.441 | 0.096 | 87.873 | < .001 |
|  |  | Valence | 0.749 | 0.072 | 59.587 | < .001 |
|  |  | Required action x cue valence | 0.019 | 0.060 | 0.093 | .760 |
| 2 | RT | Required action | -0.109 | 0.019 | 27.494 | < .001 |
|  |  | Valence | -0.191 | 0.019 | 59.204 | < .001 |
|  |  | Required action x cue valence | 0.031 | 0.015 | 4.384 | .036 |
| 3 | Accuracy | Congruency | 0.600 | 0.052 | 67.867 | < .001 |
|  |  | Stakes | -0.026 | 0.020 | 1.430 | .232 |
|  |  | Congruency x Stakes | -0.007 | 0.020 | 0.094 | .759 |
| 4 | RT | Congruency | -0.131 | 0.015 | 49.546 | < .001 |
|  |  | Stakes | 0.072 | 0.010 | 33.702 | < .001 |
|  |  | Congruency x Stakes | -0.019 | 0.010 | 3.856 | .049 |

## Supplemental Material S02: Overview results mixed-effects regression models on reaction times on all trials

**Table S02.**
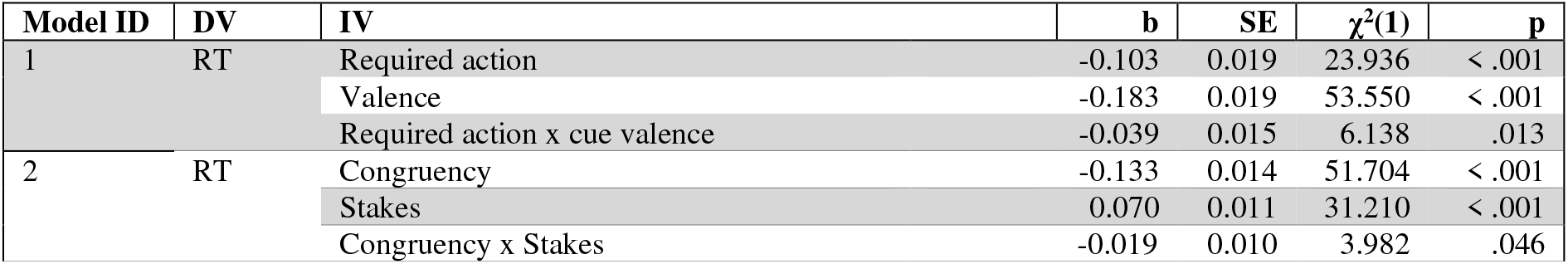
Overview of RT regression models from *N* = 54 participants when including all trials (also those with RTs < 0.3 sec.).

## Supplemental Material S03: Overview results mixed-effects regression models including additional participant

**Table S03.**
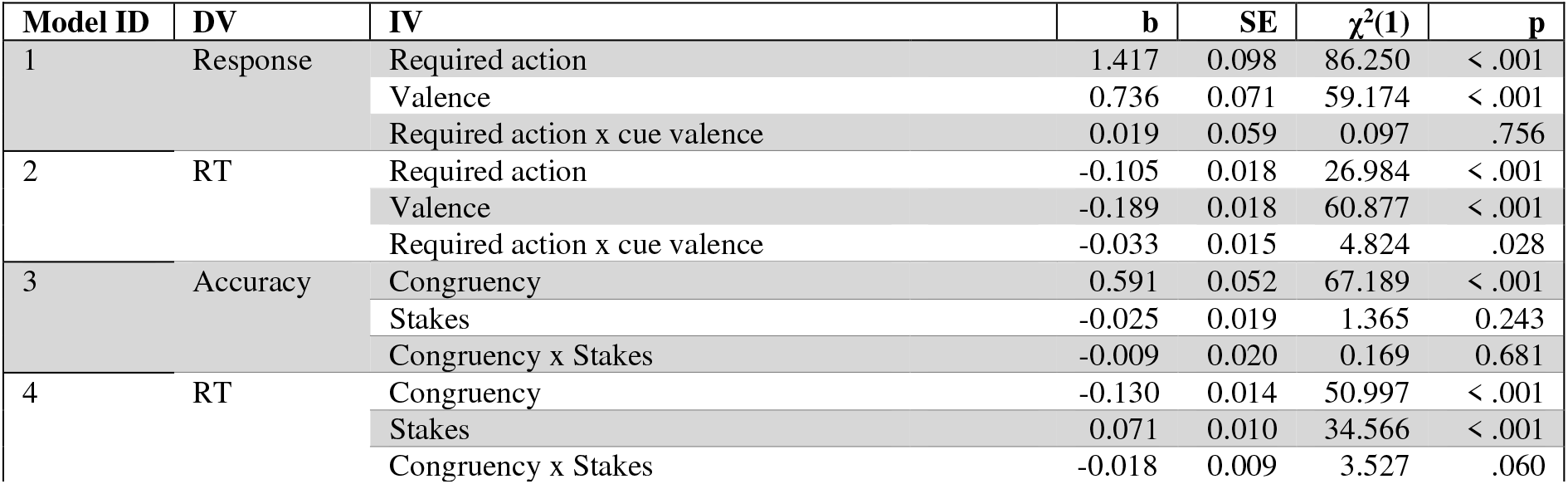
Overview of all regression models when including data from all *N* = 55 participants (also the one participant excluded from analyses reported in the main text for not performing above chance level).

## Supplemental Material S04: Overview response, accuracy, and RT means and standard deviations per condition

**Table S04.** *Means and standard deviations of Go/NoGo responses across participants per required action x valence condition*.

| Req. Act. | Go | Go | NoGo | NoGo |
| --- | --- | --- | --- | --- |
| Valence | Win | Avoid | Win | Avoid |
| Mean | 0.888 | 0.771 | 0.475 | 0.194 |
| SD | 0.140 | 0.111 | 0.261 | 0.107 |

**Table S05.** *Means and standard deviations of Go/NoGo responses across participants per required action x valence x stakes condition*.

| Req. Act. | Go | Go | Go | Go | NoGo | NoGo | NoGo | NoGo |
| --- | --- | --- | --- | --- | --- | --- | --- | --- |
| Valence | Win | Win | Avoid | Avoid | Win | Win | Avoid | Avoid |
| Stakes | High | Low | High | Low | High | Low | High | Low |
| Mean | 0.883 | 0.893 | 0.767 | 0.774 | 0.477 | 0.474 | 0.200 | 0.187 |
| SD | 0.151 | 0.138 | 0.125 | 0.114 | 0.253 | 0.276 | 0.116 | 0.109 |

**Table S06.**
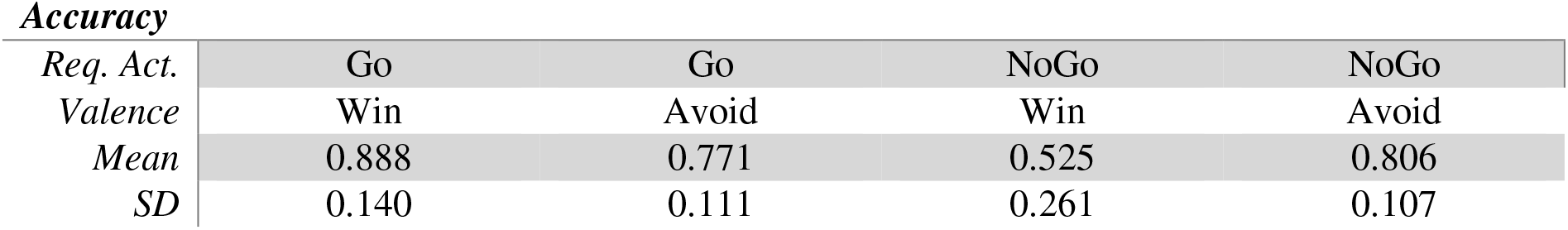
*Means and standard deviations of accuracy across participants per required action x valence condition*.

| Req. Act. | Go | Go | NoGo | NoGo |
| --- | --- | --- | --- | --- |
| Valence | Win | Avoid | Win | Avoid |
| Mean | 0.888 | 0.771 | 0.525 | 0.806 |
| SD | 0.140 | 0.111 | 0.261 | 0.107 |

**Table S07.** *Means and standard deviations of accuracy across participants per required action x valence x stakes condition*.

| Req. Act. | Go | Go | Go | Go | NoGo | NoGo | NoGo | NoGo |
| --- | --- | --- | --- | --- | --- | --- | --- | --- |
| Valence | Win | Win | Avoid | Avoid | Win | Win | Avoid | Avoid |
| Stakes | High | Low | High | Low | High | Low | High | Low |
| Mean | 0.883 | 0.893 | 0.767 | 0.774 | 0.523 | 0.526 | 0.800 | 0.813 |
| SD | 0.151 | 0.138 | 0.125 | 0.114 | 0.253 | 0.276 | 0.116 | 0.109 |

**Table S08.** *Means and standard deviations of reaction times across participants per required action x valence condition*.

| Req. Act. | Go | Go | NoGo | NoGo |
| --- | --- | --- | --- | --- |
| Valence | Win | Avoid | Win | Avoid |
| Mean | 0.578 | 0.660 | 0.641 | 0.687 |
| SD | 0.059 | 0.062 | 0.085 | 0.098 |

**Table S09.** *Means and standard deviations of reaction times across participants per required action x valence x stakes condition*.

| Req. Act. | Go | Go | Go | Go | NoGo | NoGo | NoGo | NoGo |
| --- | --- | --- | --- | --- | --- | --- | --- | --- |
| Valence | Win | Win | Avoid | Avoid | Win | Win | Avoid | Avoid |
| Stakes | High | Low | High | Low | High | Low | High | Low |
| Mean | 0.585 | 0.570 | 0.675 | 0.645 | 0.660 | 0.620 | 0.695 | 0.681 |
| SD | 0.064 | 0.063 | 0.072 | 0.063 | 0.098 | 0.099 | 0.117 | 0.122 |

## Supplemental Material S05: Correlations with questionnaires

In line with the exploratory analysis plans in mentioned in our pre-registration, we extracted the per-participant coefficients (fixed plus random effects) for (a) the effect of cue valence on responses (Pavlovian bias), (b) the effect of stakes on accuracy, (c) the effect of valence on RTs (Pavlovian bias), and (d) the effect of stakes on RTs. We then computed correlations of these coefficients with forward memory span (Fitzpatrick et al., 2015), backwards memory span, the non-planning subscale of the Barratt Impulsiveness Scale (Patton, Stanford, & Barratt, 1995), and the neuroticism subscale of the neuroticism sub-scale of the Big Five Aspects Scales (DeYoung, Quilty, & Peterson, 2007). One might plausibly hypothesize that impulsivity is related to the Pavlovian bias since many impulsive behaviors can be conceptualized as automatic, cue-triggered behaviors. Hence, individuals high on impulsivity might show stronger Pavlovian biases in responses and reaction times. Furthermore, one might hypothesize that the phenomenon of choking under pressure arises from rumination and worrying, which is typically increased in individuals scoring high on neuroticism (DeCaro, Thomas, Albert, & Beilock, 2011). Also, the effects of rumination on performance might be stronger in individuals with a low working memory score (Beilock & Carr, 2005; Bijleveld & Veling, 2014; DeCaro et al., 2011). Hence, individuals high on neuroticism and/or low on working memory span might show stronger effects of stakes on behavior.

See Figures S01 and S02 for scatterplots of all bivariate associations. None of the correlations were significant, providing no evidence for the strength of the Pavlovian bias or the effect of stakes on responses and RTs being related to either working memory span, impulsivity, or neuroticism.

**Figure S01.**
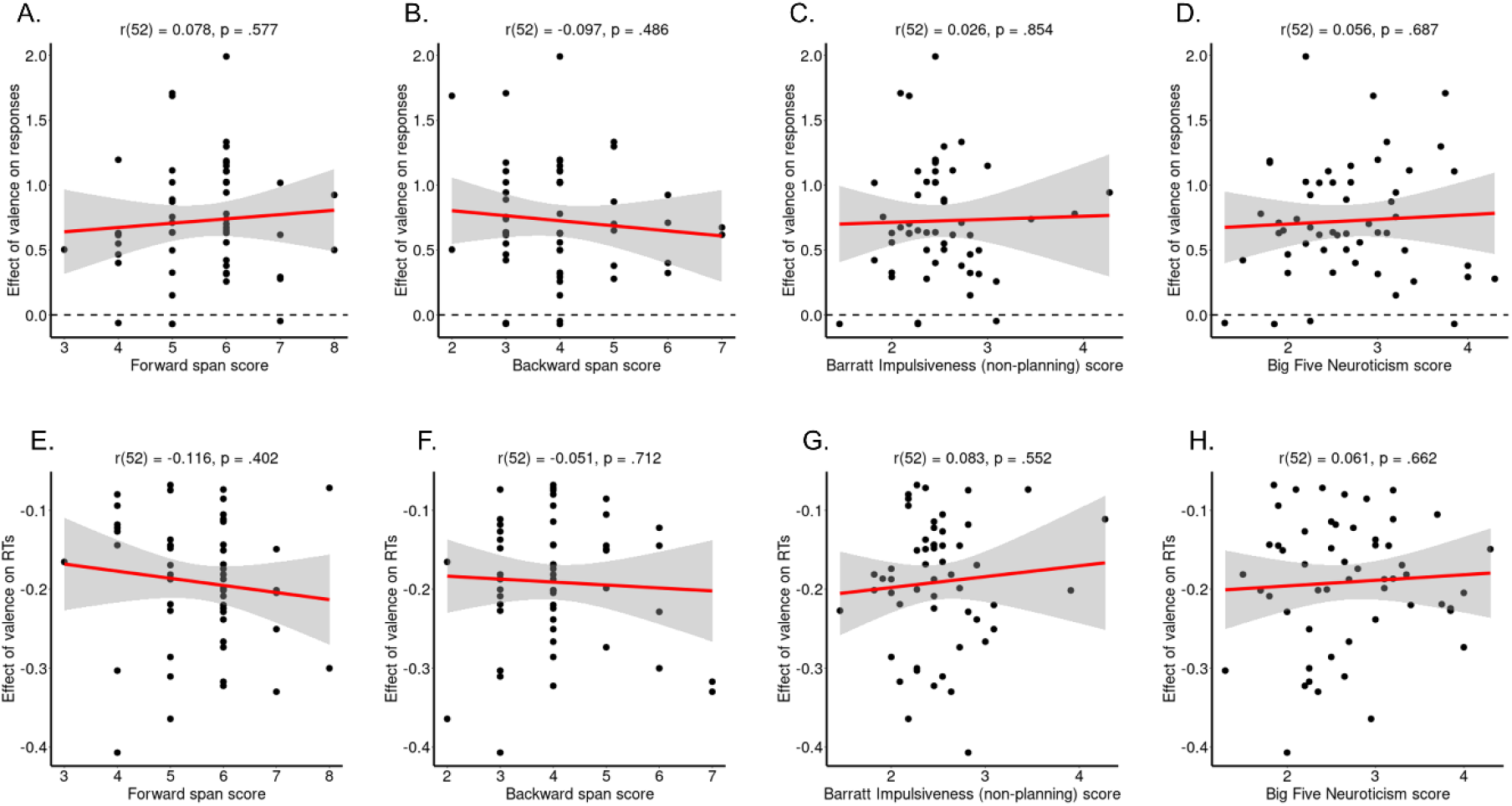
*Association of memory performance, impulsivity, and neuroticism with the valence and stakes effects on responses*. Correlations between the effect of valence on responses (**A–D**), reflecting Pavlovian biases, and the effect of stakes on accuracy (**E–H**) with (**A/F**) forward working memory span, (**B/F**) backwards working memory span, (**C/G**) impulsivity (Barratt Impulsiveness Scale, non-planning subscale) and (**D/H**) neuroticism. Black dots represent per-participant scores, the red line the best-fitting regression line, they grey shade the 95%-confidence interval. None of the displayed correlations is significant at α = .05.

**Figure S02.**
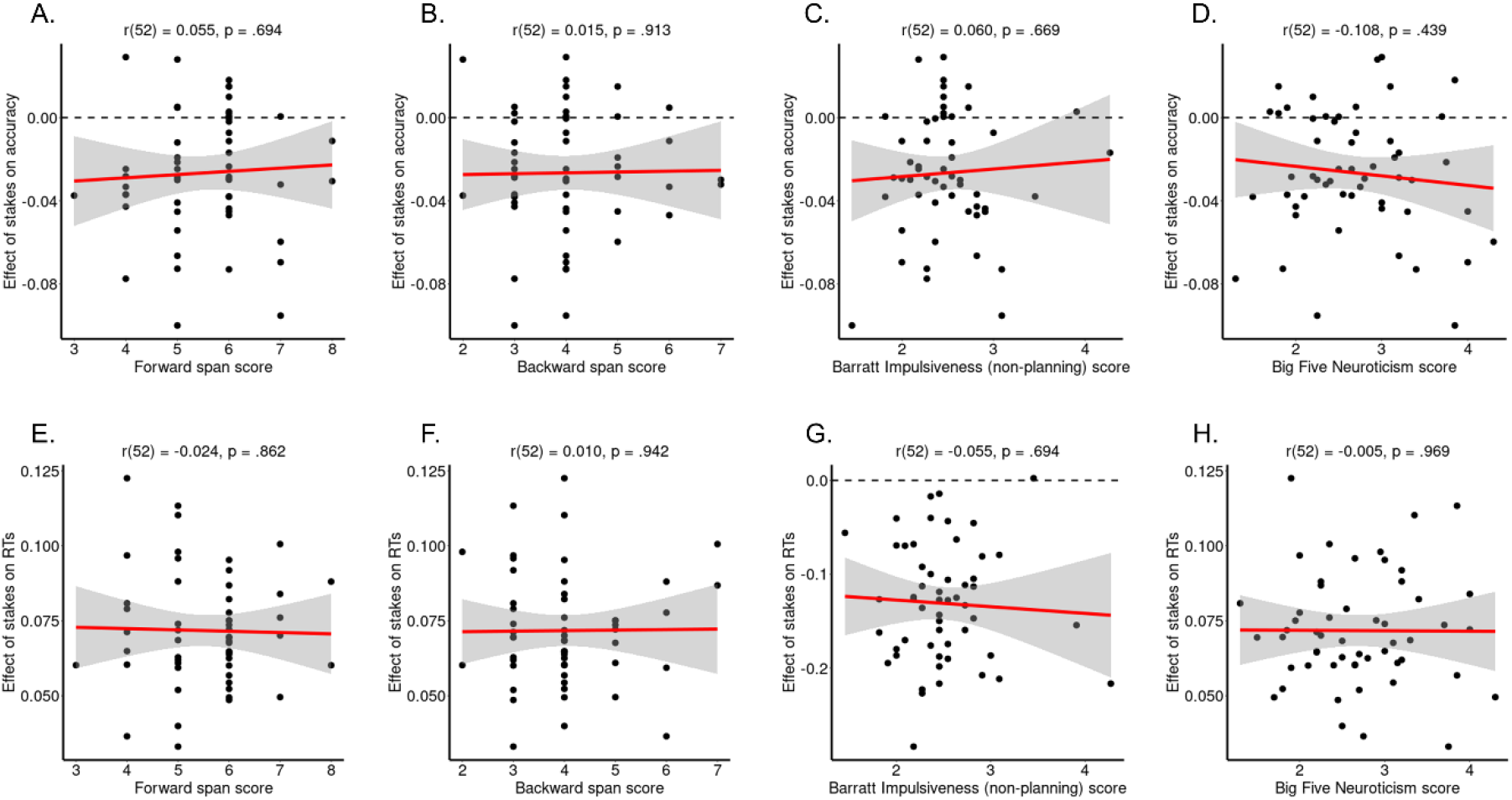
*Association of memory performance, impulsivity, and neuroticism with the valence and stakes effects on RTs*. Correlations between the effect of valence on RTs (**A–D**), reflecting Pavlovian biases, and the effect of stakes on RTs (**E–H**) with (**A/F**) forward working memory span, (**B/F**) backwards working memory span, (**C/G**) impulsivity (Barratt Impulsiveness Scale, non-planning subscale) and (**D/H**) neuroticism. Black dots represent per-participant scores, the red line the best-fitting regression line, they grey shade the 95%-confidence interval. None of the displayed correlations is significant at α = .05.

## Supplemental Material S06: Effect of stakes on RTs over time

In the results in the main text, we report linear associations between time on task (cue repetition, trial number with blocks, trial number across blocks) and reaction time. All associations were non-significant. A more sensitive approach to detect possible non-linear changes over time are so called additive models, which model a time series as a mixture of smooth functions (i.e., thin plate regression splines) for each condition, and allow to test whether (a) a given time series is significant different from a flat line, and (b) whether the time series of different conditions are significantly different from each other (Baayen et al., 2017; Wood, 2017). A smooth function regularizes a raw times series and suppresses high-frequency (i.e., trial-by-trial) noise. Furthermore, it allows for non-zero auto-correlation between residuals, which are assumed to be zero in linear models.

In order to test whether the effect of task conditions of stakes on RTs changed over time, we fit three generalized additive mixed-effects models with the z-standardized trial-by-trial RT as dependent variable, modelled as an effect of cue repetition (1–20) with separate time series for (a) each cue condition (Go-to-Win, Go-to-Avoid, NoGo-to-Win, NoGo-to-Avoid), (b) for each stakes condition (high, low), or (c) the interaction between congruency (congruent, incongruent) and stakes (high, low). We modeled the time course of cue repetition as a factor smooth (which has a similar, but potentially non-linear effect as adding a random intercept and a random slope) for each participant for each block, allowing for the possibility that condition differences were different in different participants in different blocks (equivalent to a full random-effects structure). We used a scaled *t*-distribution instead of a Gaussian distribution for the RT variable as it led to lower AIC values. We also investigated whether fit further improved by adding an AR(1) auto-regressive model, which was not the case. For all fitted models, We visually checked that residuals were approximately normally distributed using quantile-quantile plots and whether auto-correlation was near zero using auto- correlation plots (van Rij et al., 2019).

The model testing for differences between cue conditions suggested that RTs overall significantly decreased over time in all conditions (see Table S10; Fig. S03A). Further, RTs started to differ between cue conditions from repetition 1 or 2 onwards (see Table S11). Overall, RTs were faster for responses to Win than Avoid cues and faster for (correct) responses to Go cues than (incorrect) responses to NoGo cues. Overall, RT differences between conditions persisted throughout the block.

The model testing for differences between stakes levels suggested again that RTs overall significantly decreased over time in both conditions (Table S10; Fig. S03B). Furthermore, throughout the block, RTs were slower for responses on high-stakes trials than for responses on low-stakes trials (Table S11). This difference persisted throughout the block.

Finally, the model testing for differences between congruency conditions and stakes levels found again a significant decrease in RTs over time (Table S10; Fig. S03C). RTs were slower for responses to incongruent than to congruent cues, and slower on high-stakes trials than on low-stakes trials. Importantly, RTs were slower on high-stakes trials compared to low-stakes trials both for congruent and for incongruent cues, similarly, although this differences tended to be bigger for incongruent trials. These differences persisted throughout the task.

In sum, these results show that condition differences and differences between stakes in RTs emerge on the very first trials (cue repetitions) of a task and persist until the end of a block, with little change in these condition differences.

**Figure S03.**
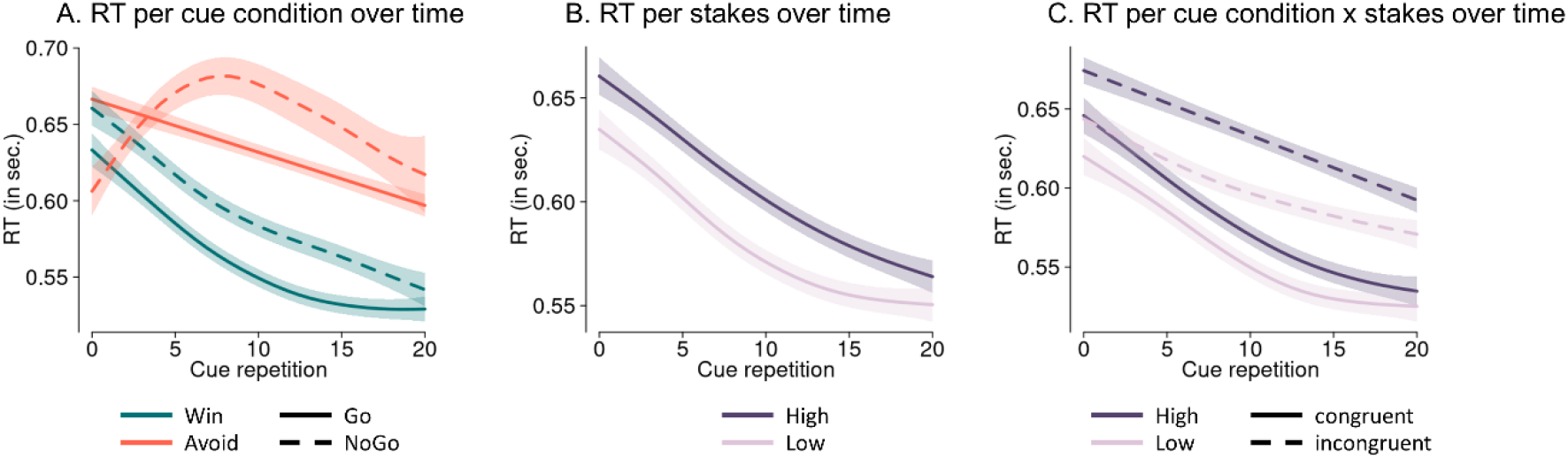
*Time course of RTs over cue repetitions within a block as predicted by a generalized additive mixed-effects model, separated by conditions*. Overall, RTs speed up over time. **A.** Differences between cue conditions as predicted by the fit of a . RTs are significantly faster for responses to Win than responses to Avoid cues, and faster for (correct) responses to Go cues than (incorrect) responses to NoGo cues throughout a block. **B**. Differences between stakes levels. RTs are significantly slower on high-stakes trials compared to low-stakes trials throughout a block. **C**. Differences between stakes levels separately per condition. Both for congruent and incongruent cues, RTs on high-stakes trials are significantly slower than RTs on low-stakes trials. This difference tends to be larger for incongruent cues.

**Table S10.** *Results from generalized additive mixed models (GAMMs) with separate smooth per condition.* The parametric term reflects a linear change in time, while the smooth terms reflects any non-linear changes. Both add up to the total term.

| Model | Parametric coefficient<br>(Linear change within condition) | Smooth<br>(non-linear change within each condition) |
| --- | --- | --- |
| **Cue conditions:** |  |  |
| Go-to-Win | $t(3, 0.103) = 115.249, p < .001$ | $F(3.094, 3.788) = 27.370, p < .001$ |
| Go-to-Avoid | $t(3, 0.103) = 22.778, p < .001$ | $F(1.000, 1.000) = 35.715, p < .001$ |
| NoGo-to-Win | $t(3, 0.103) = 8.036, p < .001$ | $F(2.497, 3.061) = 26.894, p < .001$ |
| NoGo-toAvoid | $t(3, 0.103) = 12.887, p < .001$ | $F(2.963, 3.629) = 1.530, p = .107$ |
| **Stakes:** |  |  |
| High | $t(3, 0.107) = 122.148, p < .001$ | $F(2.266, 2.746) = 35.926, p < .001$ |
| Low | $t(3, 0.107) = -9.346, p < .001$ | $F(2.857, 3.478) = 24.505, p < .001$ |
| **Congruency x Stakes:** |  |  |
| Congruent /high | $t(3, 0.104) = 108.940, p < .001$ | $F(2.426, 2.973) = 30.546, p < .001$ |
| Congruent/ low | $t(3, 0.104) = -4.957, p < .001$ | $F(2.679, 3.284) = 22.242, p < .001$ |
| Incongruent/ high | $t(3, 0.104) = 15.148, p < .001$ | $F(1.000, 1.000) = 44.597, p < .001$ |
| Incongruent/ low | $t(3, 0.104) = 6.496, p < .001$ | $F(1.947, 2.381) = 15.505, p < .001$ |

**Table S11.** *Results from generalized additive mixed models (GAMMs) with difference smooths between two conditions.* The parametric term reflects a linear difference between conditions, while the smooth terms reflects any non-linear difference. Both add up to the total term. The time window of significant condition differences is automatically returned by the model.

| *Model* | **Parametric coefficient**<br>**(Intercept difference)** | **Smooth**<br>**(non-linear differences)** | **Windows of**<br>**significant**<br>**differences** |
| --- | --- | --- | --- |
| ***Cue conditions:*** |  |  |  |
| *G2W – G2A* | $t(3, 0.096) = 23.320, p < .001$ | $F(3.779, 4.618) = 5.052, p < .001$ | 1 – 20 |
| *G2W – NG2W* | $t(3, 0.081) = 8.383, p < .001$ | $F(1.000, 1.000) = 0.606, p = .436$ | 0 – 20 |
| *G2W – NG2A* | $t(3, 0.081) = 12.400, p < .001$ | $F(3.255, 3.933) = 14.710, p < .001$ | 2 – 20 |
| *G2A – NG2W* | $t(3, 0.108) = -9.870, p < .001$ | $F(2.587, 3.168) = 5.080, p = .001$ | 2 – 20 |
| *G2A – NG2A* | $t(3, 0.112) = 3.234, p = .001$ | $F(2.878, 3.476) = 7.412, p < .001$ | 0 – 2, 5 – 16 |
| *NG2W – NG2A* | $t(3, 0.098) = 6.939, p < .001$ | $F(3.376, 4.042) = 11.760, p < .001$ | 0 – 1, 3 – 20 |
| ***Stakes:*** |  |  |  |
| *High – Low* | $t(3, 0.107) = -9.317, p < .001$ | $F(1.424, 1.706) = 1.715, p = .278$ | 0 – 20 |
| ***Congruency x Stakes:*** |  |  |  |
| *Cong/High – Cong/Low* | $t(3, 0.081) = -4.997, p < .001$ | $F(1.000, 1.000) = 0.039, p = .844$ | 0 – 20 |
| *Incong/High – Incong/Low* | $t(3, 0.108) = -8.337, p < .001$ | $F(1.000, 1.000) = 0.369, p = .543$ | 0 – 20 |
| *Cong/High – Incong/High* | $t(3, 0.999) = 15.430, p < .001$ | $F(1.711, 2.085) = 2.757, p = .061$ | 0 – 20 |
| *Cong/Low – Incong/Low* | $t(3, 0.102) = 11.470, p < .001$ | $F(1.000, 1.000) = 5.196, p = .023$ | 0 – 20 |

## Supplemental Material S07: Reinforcement-learning drift diffusion models

**Table S12.** *Mean [25^th^ percentile, 75^th^ percentile] of the posterior densities of group-level parameters.* α = decision threshold, τ = non-decision time, β = starting point bias, δ = drift rate, ε = learning rate. WAIC and LOO-IC are reported as measures of model fit, with smaller values indicating a better fit.

|  | M01 | M02 | M03 | M04 | M05 | M06 | M07 | M08 | M09 | M10 | M11 | M12 |
| --- | --- | --- | --- | --- | --- | --- | --- | --- | --- | --- | --- | --- |
| WAIC | 14501 | 9284 | 8011 | 7029 | 7023 | 6848 | 7082 | 6996 | 6821 | 7025 | 6843 | 6682 |
| LOO-IC | 14365 | 8970 | 7611 | 6734 | 6656 | 6512 | 6722 | 6706 | 6420 | 6646 | 6494 | 6278 |
| $\alpha$ | 2.011<br>[1.97,<br>2.02] | 1.377<br>[1.372,<br>1.404] | 1.466<br>[1.449,<br>1.482] | 1.442<br>[1.428,<br>1.456] | | 1.409<br>[1.397,<br>1.421] | 1.430<br>[1.417,<br>1.444] | 1.444<br>[1.430,<br>1.457] | | | 1.408<br>[1.396,<br>1.421] | 1.379<br>[1.367,<br>1.390] |
| $\alpha_{Low}$ | | | | | 1.406<br>[1.393,<br>1.420] | | | | 1.375<br>[1.361,<br>1.389] | 1.410<br>[1.397,<br>1.424] | | |
| $\alpha_{High}$ | | | | | 1.479<br>[1.466,<br>1.492] | | | | 1.429<br>[1.414,<br>1.444] | 1.475<br>[1.462,<br>1.488] | | |
| $\tau$ | 0.128<br>[0.119,<br>0.136] | 0.234<br>[0.226,<br>0.241] | 0.228<br>[0.220,<br>0.236] | 0.232<br>[0.224,<br>0.239] | 0.233<br>[0.226,<br>0.240] | | 0.234<br>[0.227,<br>0.241] | 0.231<br>[0.224,<br>0.239] | | 0.233<br>[0.225,<br>0.240] | | |
| $\tau_{Low}$ | | | | | | 0.237<br>[0.230,<br>0.245] | | | 0.243<br>[0.236,<br>0.250] | | 0.238<br>[0.231,<br>0.245] | 0.244<br>[0.237,<br>0.251] |
| $\tau_{High}$ | | | | | | 0.249<br>[0.242,<br>0.256] | | | 0.247<br>[0.240,<br>0.254] | | 0.249<br>[0.241,<br>0.256] | |
| $\tau_{High/Cong}$ | | | | | | | | | | | | 0.266<br>[0.260,<br>0.273] |
| $\tau_{High/Incong}$ | | | | | | | | | | | | 0.264<br>[0.256,<br>0.271] |
| $\beta$ | 0.061<br>[0.058,<br>0.063] | 0.259<br>[0.251,<br>0.267] | | 0.251<br>[0.244,<br>0.258] | 0.250<br>[0.243,<br>0.257] | 0.264<br>[0.257,<br>0.270] | | 0.249<br>[0.243,<br>0.256] | 0.266<br>[0.259,<br>0.273] | 0.250<br>[0.243,<br>0.256] | 0.264<br>[0.257,<br>0.271] | 0.277<br>[0.270,<br>0.284] |
| $\beta_{Win}$ | | | 0.318<br>[0.308,<br>0.328] | | | | | | | | | |
| $\beta_{Avoid}$ | | | 0.167<br>[0.161,<br>0.172] | | | | | | | | | |
| $\beta_{Low}$ | | | | | | | 0.268<br>[0.262,<br>0.274] | | | | | |
| $\beta_{High}$ | | | | | | | 0.247<br>[0.241,<br>0.253] | | | | | |
| $\delta_{Int}$ | 3.617<br>[3.558,<br>3.675] | 1.358<br>[1.310,<br>1.407] | 1.542<br>[1.483,<br>1.602] | | | | | | | | | |
| $\delta_{Win}$ | | | | 2.086<br>[2.018,<br>2.152] | 2.100<br>[2.032,<br>2.167] | 2.037<br>[1.890,<br>2.105] | 2.041<br>[1.971,<br>2.112] | 2.159<br>[2.091,<br>2.228] | 2.026<br>[1.957,<br>2.094] | 2.130<br>[2.061,<br>2.199] | 2.074<br>[2.005,<br>2.142] | 1.981<br>[1.910,<br>2.053] |
| $\delta_{Avoid}$ | | | | 0.796<br>[0.757,<br>0.834] | 0.803<br>[0.765,<br>0.841] | 0.736<br>[0.698,<br>0.774] | 0.763<br>[0.726,<br>0.799] | 0.867<br>[0.827,<br>0.908] | 0.727<br>[0.691,<br>0.764] | 0.831<br>[0.791,<br>0.871] | 0.774<br>[0.736,<br>0.813] | 0.684<br>[0.645,<br>0.723] |
| $\delta_{Slope}$ | | 6.823<br>[6.354,<br>7.267] | 6.283<br>[5.872,<br>6.681] | 6.093<br>[5.718,<br>6.446] | 6.149<br>[5.773,<br>6.508] | 6.191<br>[5.810,<br>6.550] | 6.102<br>[5.734,<br>6.458] | 6.109<br>[5.741,<br>6.464] | 6.218<br>[5.834,<br>6.590] | 6.151<br>[5.777,<br>6.510] | 6.219<br>[5.834,<br>6.586] | 6.273<br>[5.896,<br>6.633] |
| $\delta_{High}$ | | | | | | | | -0.128<br>[-0.148,<br>-0.108] | | -0.054<br>[-0.076,<br>-0.033] | -0.075<br>[-0.096,<br>-0.054] | |
| $\varepsilon$ | | 0.100<br>[0.088,<br>0.110] | 0.102<br>[0.092,<br>0.112] | 0.121<br>[0.110,<br>0.131] | 0.121<br>[0.109,<br>0.131] | 0.120<br>[0.108,<br>0.130] | 0.120<br>[0.109,<br>0.130] | 0.122<br>[0.110,<br>0.132] | 0.120<br>[0.109,<br>0.130] | 0.121<br>[0.109,<br>0.131] | 0.120<br>[0.109,<br>0.131] | 0.119<br>[0.108,<br>0.129] |

**Figure S04.**
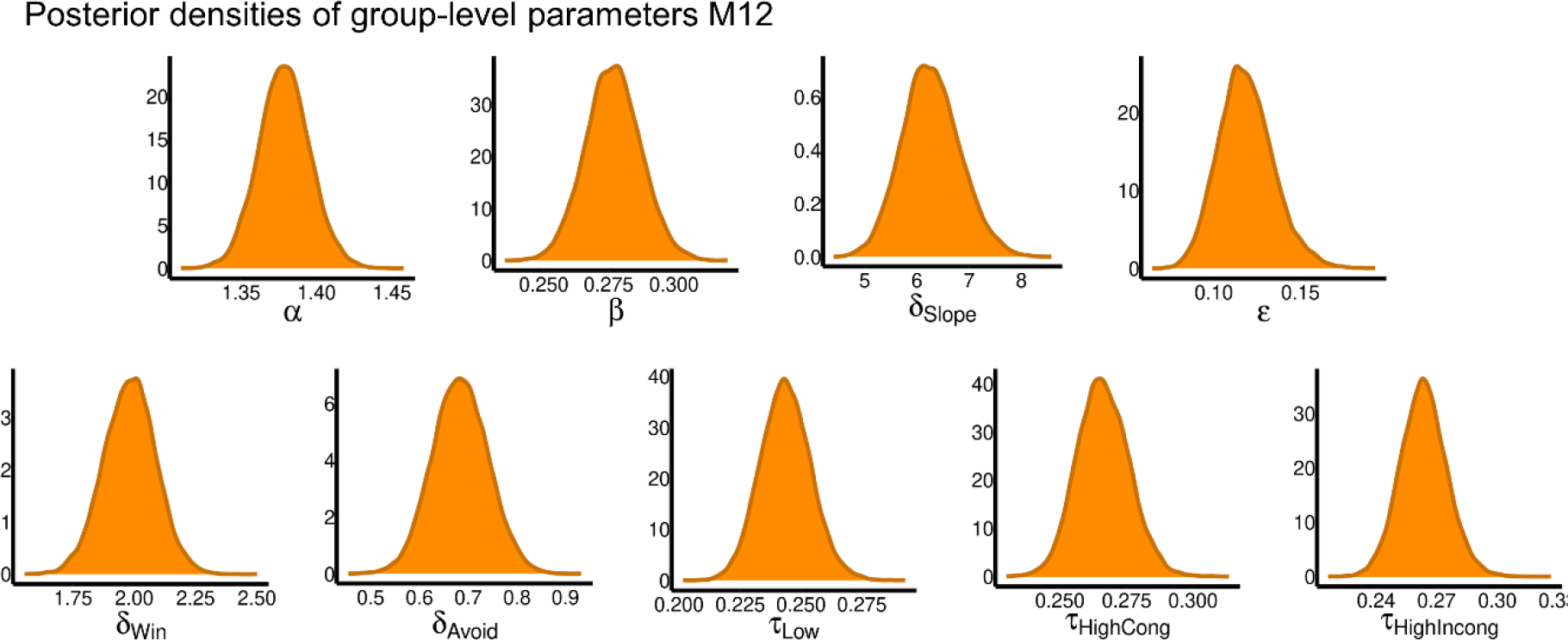
*Posterior densities of the group-level parameters of the winning model M12*. α = decision threshold, τ = non-decision time, β = starting point bias, δ = drift rate, ε = learning rate.

**Figure S05.**
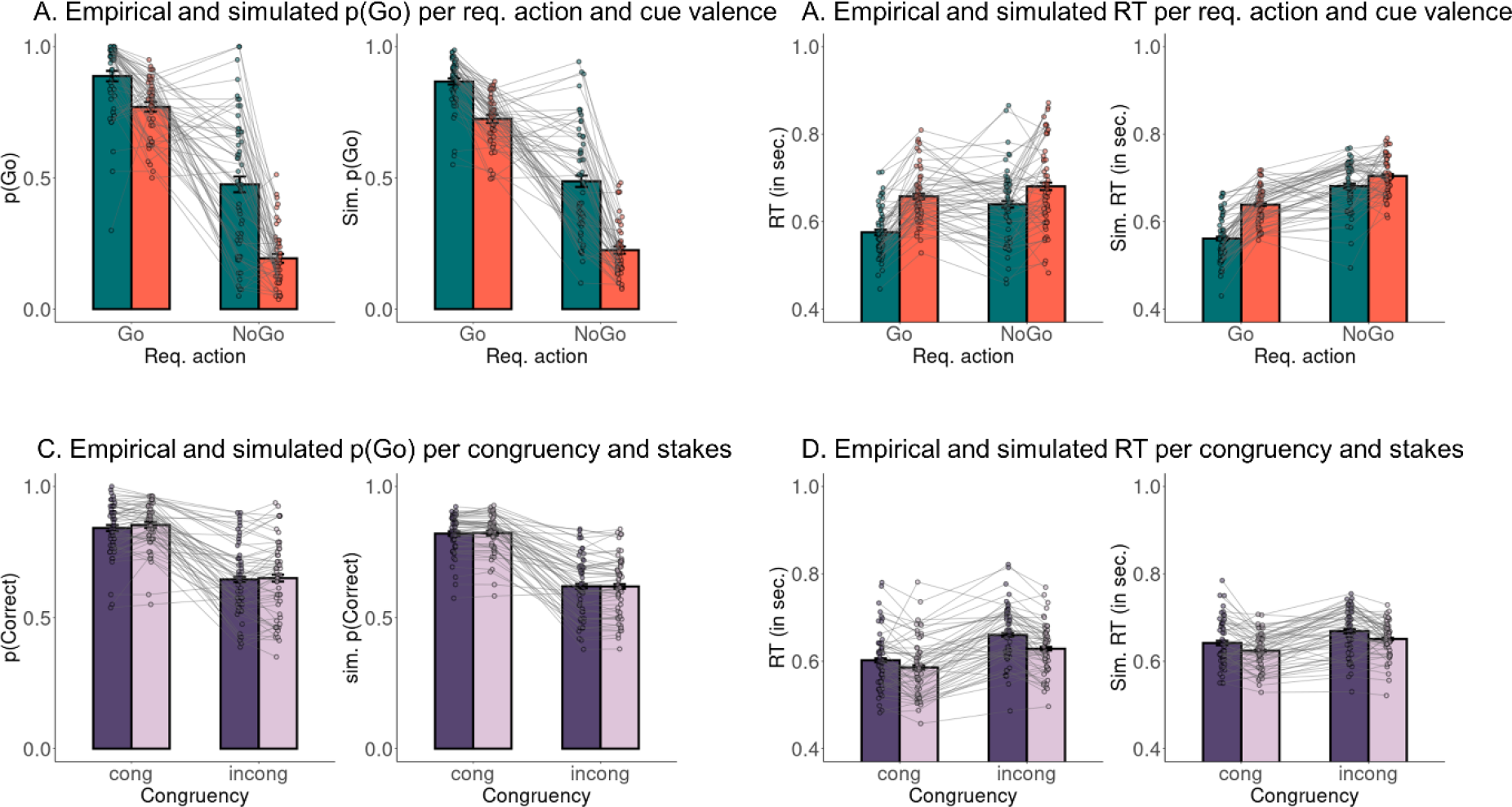
*Posterior predictive checks for data simulated from the winning model M12*. **A.** Both in empirical data (left panel) and data simulated from the winning model M12 (right panel), (simulated) participants performed more Go responses to Go than NoGo cues (learning) and more Go responses to Win than Avoid cues (Pavlovian bias). Simulated data matched the empirical data pattern. **B**. Both in empirical and simulated data, (simulated) participants showed faster responses to Go than NoGo cues and to Win than Avoid cues. Simulated data matched the empirical data pattern. **C**. Both in empirical and simulated data, (simulated) participants performed more accurately for congruent than incongruent cues, with no difference between high and low stakes. **D**. Both in empirical and simulated data, (simulated) participants performed faster for congruent than incongruent cues and under low compared to high stakes. In empirical participants, the stakes effect was stronger for incongruent than congruent cues, but this difference was somewhat underestimated by the winning model M12.

**Figure S06.**
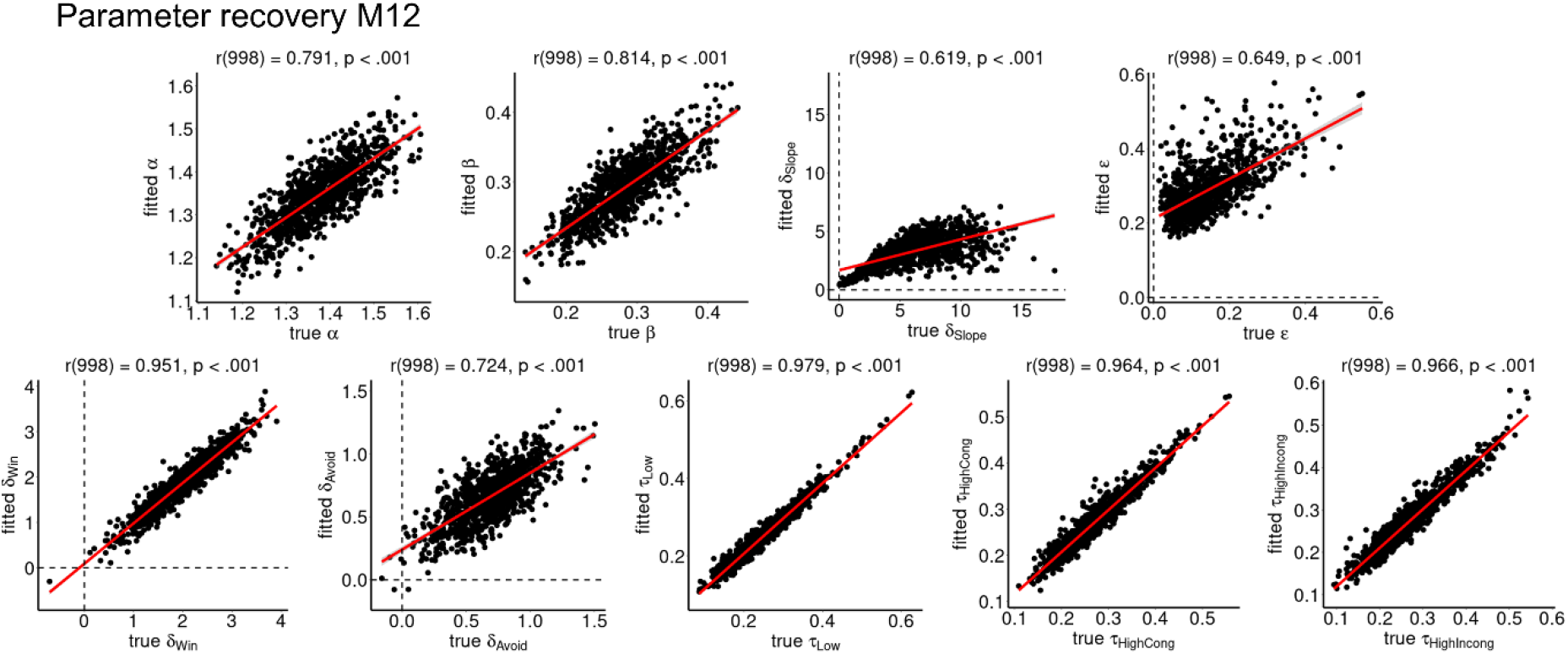
*Parameter recovery results for the winning model M12*. The correlation between generative and fitted parameters is overall very high. Recovery is overall very high. It is least optimal (but still strongly significant) for δSlope and ε, which trade off against each other (see Fig. 4D main text). α = decision threshold, τ = non-decision time, β = starting point bias, δ = drift rate, ε = learning rate.

**Figure S07.**
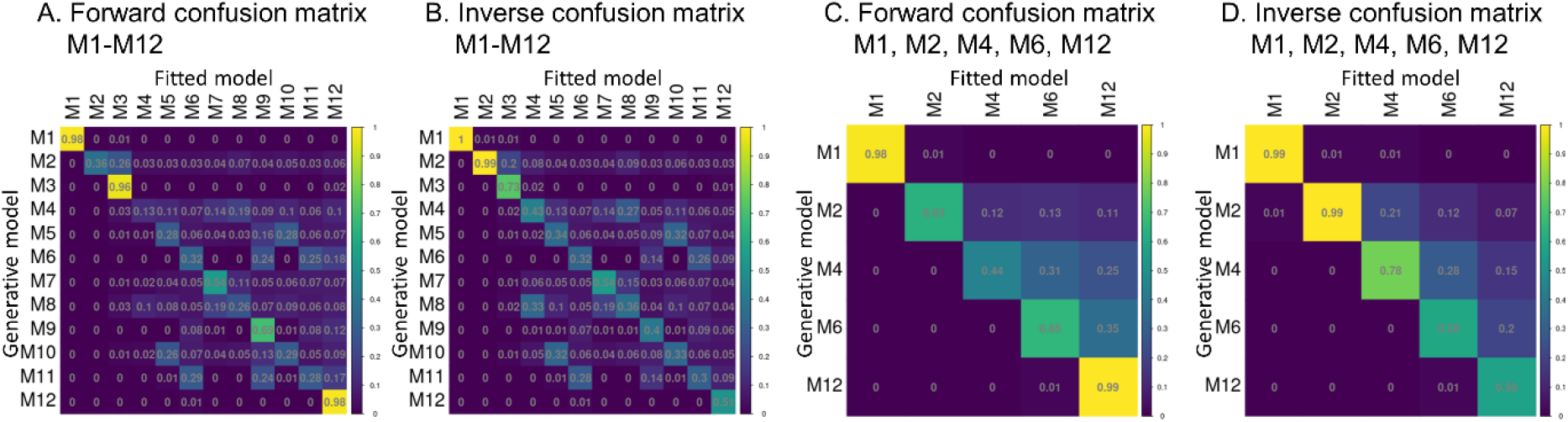
*Forward and inverse confusion matrices from model recovery of all models and of nested sub-versions of the winning model M12*. **A**. The forward confusion matrix displays the conditional probabilities that model Y is the best fitting model (columns) if model X (rows) is the underlying generative model used to simulate a given data set (identical to Fig. 4E main text). Rows sum to 100%. On-diagonal probabilities indicate the probability of reidentifying the generative model. All on-diagonal probabilities are significantly above chance (range 0.13–0.98; 95th percentile of permutation null distribution: p = 0.10). Especially recovery for M12 is exceptionally high (98%). **B.** The inverse confusion matrix displays the conditional probabilities that model X is the generative model (rows) if model Y (rows) is the best fitting model for a given data set. Columns sum to 100%. On-diagonal probabilities indicate the probability of reidentifying the generative model. All on-diagonal probabilities are significantly above chance (range 0.30–1.00; 95th percentile of permutation null distribution: p = 0.10). **C**. Forward confusion matrix only for the five models that are nested sub-versions of M12 (i.e., M1, M2, M4, M6, M12). Recovery is overall much higher (range 0.44–0.99; 95^th^ percentile of permutation null distribution: p = 0.22). **D**. Inverse confusion matrix only for the five models that are nested sub-versions of M12. Recovery is overall much higher (range 0.58–0.99; 95^th^ percentile of permutation null distribution: p = 0.22).

## Notes

### Competing Interest Statement

The authors have declared no competing interest.

### Summary of Updates

1.Major rewriting of the Introduction section, so that we now: a.explicitly define Pavlovian biases b.motivate why Pavlovian biases should become stronger under high stakes 2.Extension of the Results section, so that we now: a.motivate and explain reinforcement-learning drift-diffusion models (RL-DDMs) and their particular implementation for our task b.document more extensively the different models and model comparison steps 3.Major rewriting of the Discussion section, so that we now: a.explain how cognitive control (specifically response slowing) can occur in situa-tions where it is irrelevant for task performance and thus used inefficiently b.discuss whether stakes might lead both to bias amplification and heightened cog-nitive control recruitment, with both effects cancelling each other out c.explain the potential mechanisms underlying positive conditioned suppression d.discuss more explicitly how serotonin might play a role in positive conditioned suppression

